# Spontaneous Phase Separation Enables Rapid, Polymerization-Free Fabrication of Dissolvable Hydrogels

**DOI:** 10.64898/2026.01.10.698843

**Authors:** Namrata Priyadarshinee, Vidhi Saxena, Aniruddha Kambekar, Gaurav Chauhan, Karthik Pushpavanam

## Abstract

Hydrogels are cross-linked polymeric networks with wide applications in drug delivery, tissue engineering, biosensing, and environmental remediation. These hydrogels additionally host living cells, small molecules and biological propagules, which further expand the applications of these materials. However, most if not all fabrication methods require covalent modifications. In this work, for the first time, we demonstrate that polymer mixtures can access an additional material state beyond the conventionally described homogeneous and two-phase regimes. By deliberately selecting polymers with a known propensity to phase separate and formulating compositions far from the binodal boundary, the system transitions directly into a mechanically stable hydrogel. We demonstrate this technique using a model system of poly (ethylene glycol) (PEG) and dextran (DEX). We have systematically characterized the hydrogels through FTIR, MALDI-TOF to discern the molecular compositions of the hydrogels. We also modulate the optical transparency of these hydrogels by varying the molecular weight of the polymers. These experimental findings are supplemented with coarse grained (CG) simulation insights to investigate the mechanistic origins of phase separation propensity with varying molecular weights of dextran. We utilized coexisting densities in the two phases using CG simulations to predict the role of dextran molecular weight on the partitioning of PEG and DEX in the two phases. Finally, we exploit the fabricated hydrogel’s ability to encapsulate live cells, antibiotics and plant seeds. We anticipate that this ATPS-based fabrication technique will provides a scalable, crosslinker-free route to multifunctional hydrogels enabling advanced applications in drug delivery and responsive materials.

## INTRODUCTION

Hydrogels are three-dimensional, hydrophilic polymer networks capable of absorbing and retaining significant amounts of water while maintaining structural integrity.^1^ The polar hydrophilic moieties of polymers capable of hydrogen bonding and ionic interactions through -NH_2_, -COOH, -CONH, -OH, -CONH_2,_ and -SO_3_H contribute to the network’s hydrophilicity.^2^ These impart hydrogels with distinctive properties, such as biodegradability and biocompatibility which underpin their extensive use across biomedical and materials science domains.^3^ Additionally, modulating the polymer composition and water content determine its optical clarity.^4^ Despite their widespread use, most if not all hydrogel fabrication techniques rely on polymerization, including bulk polymerization, cryogelation and frontal polymerization.^5, 6, 7, 8^ These processes typically involve the use of small molecule initiators and reactive crosslinkers.^9, 10, 11^ These reactive steps limit biocompatibility and constrain the encapsulation of sensitive biological cargo such as live cells or labile molecules during polymerization.^12, 13, 14^ There is still a need for a polymerization-free hydrogel fabrication strategy, which could circumvent the above-mentioned challenges.

Recently, hydrogels designed through modulating hydrogen bonding, electrostatic attraction, π-π stacking have emerged as alternatives to chemically cross-linked hydrogels.^15^ A major class of these hydrogels are through short peptide sequences, typically smaller than 20 residues, such as RADA16-I and β-hairpin MAX1. These undergo self-assembly into nanofibrous networks through ionic complementarity and hydrogen bonding, resulting in the formation of hydrogels.^16, 17, 18, 19^ These hydrogels due to their high biocompatibility have been utilized in a wide range of biomedical applications including as live cell, small molecule carriers.^19, 20^ However, comparable self-assembly strategies have not been explored for larger polymers (>10,000 Da). Hydrogels formed through non-covalent interactions between polymers could offer enhanced mechanical strength.^15^ Finally, their broad availability could ensure scalability unlike peptide-based hydrogels which typically relies on solvent-intensive solid-phase peptide synthesis.^15^

Recently, we along with another research group had demonstrated the use of aqueous two-phase systems (ATPS) to generate dual-layered hydrogels.^21^ Although effective, we still employed polyacrylamide crosslinking to generate these hydrogels. Given that ATPS emerges from a binary mixture of polymers, we posed whether pushing the polymer composition far from the binodal curve could enable hydrogel formation directly from ATPS through polymer entanglement. This would eliminate the need for chemical crosslinking. This independence from polymerization will expand the hydrogels utility, particularly for applications that require mild, biocompatible, or modular fabrication workflows.^22, 23, 24^ Motivated by this gap, we introduce a polymerization-free, one-step approach for generating dissolvable, optically clear hydrogels by using GRAS (Generally Recognized as Safe) biopolymers.

## MATERIALS AND METHODS

### Materials

Polyethylene Glycol 400 (PEG 400) for molecular biology (CAS Number: 25322-68-3), Dextran average mol wt. 9,000-11,000 (DEX 10 kDa) from *Leuconostoc mesenteroides* (CAS Number: 9004-54-0), and Dextran average mol wt. 450,000-650,000 (DEX 500 kDa) from *Leuconostoc mesenteroides* (CAS Number: 9004-54-0), Poly(2-ethl-2-oxazoline) (CAS Number: 25805-17-8) were purchased from Sigma-Aldrich. Dextran 70,000 (DEX 70 kDa) from *Leuconostoc mesenteroides* (CAS Number: 9004-54-0), and Coomassie Brilliant Blue R-250 (CBB) for molecular biology (CAS Number: 6104-59-2), Ferric chloride Anhydrous pure (CAS Number: 7705-08-0) were purchased from Sisco Research Laboratories Pvt. Ltd. Chloroform (CAS Number: 67-66-3) was purchased from Loba Chemie Pvt Ltd. Ammonium thiocyanate (CAS Number: 1762-95-4) was purchased from TCI Chemicals. Milli-Q water was used as the solvent unless otherwise mentioned.

### Generation of Binodal Curve via the Cloud Point Method

Stock solutions of 40 wt.% Polyethylene glycol (PEG 400) and 40 wt.% Dextran (DEX) with varying molecular weights (10 kDa, 70 kDa and 500 kDa) were prepared. To determine the binodal curve, PEG and DEX solutions at different concentrations were mixed and vortexed for 30 s. The cloud point was identified visually by observing the transition from a transparent to a turbid solution. The PEG and DEX concentrations corresponding to the onset of turbidity were recorded. The critical binodal points were refined by progressively narrowing the concentration range, allowing precise mapping of the binodal curve.^25^

### Generation of PEG and DEX Hydrogels

Hydrogels were synthesized by combining PEG 400 with Dextran (DEX) of varying molecular weights. Briefly, in a 0.5 ml tube, 20 µl of 100 wt.% PEG 400 and 20µl of 40 wt.% DEX of the appropriate molecular weight stock solution was mixed. The mixture was vortexed thoroughly for 3-5 s to achieve the hydrogel formation.

### Preparation of Lyophilized Hydrogel

The hydrogel was first frozen at –80 °C for 3-4 h, followed by lyophilization until complete drying.

### Colorimetric assay to determine PEG concentration

A colorimetric assay devised by Nag et al. was employed to quantify the concentration of PEG in the supernatant after the gel formation and in the milli-Q water dissolved hydrogel.^26^ In a 1.5 mL tube, 0.5 mL of ammonium ferro thiocyanate and 0.5 mL of chloroform were combined, followed by the addition of 50 µL of the sample. The mixture was vortexed vigorously for 30 min and centrifuged at 3000 rpm for 2 min. The chloroform (lower) phase containing PEG was carefully extracted, and its absorbance was measured at 510 nm using a SHIMADZU UV-1800 spectrophotometer with a 10 mm path-length cuvette. All measurements were performed in triplicate.

### Fourier Transform Infrared Spectroscopy (FTIR) Analysis

FTIR spectra were recorded using a PerkinElmer FTIR spectrophotometer across the spectral range of 4000–500 cm^−1^, to investigate the spectra of PEG 400, DEX of different molecular weights (10 kDa, 70 kDa, and 500 kDa), and their corresponding hydrogels.

### Dye Dissolution Study

3.5 mg/mL of Coomassie Brilliant Blue dye was added to the 40 wt.% DEX stock solution prior to the hydrogel formation. The hydrogel was transferred into a petri dish containing 25 mL of Milli-Q water, and its dissolution behaviour was recorded over time. The reduction in hydrogel size was quantified using ImageJ software, following a standardized image-processing protocol to determine dissolution kinetics.

### Matrix-assisted laser desorption/ionization time-of-flight mass spectrometry (MALDI-TOF MS)

PEG 400, DEX of different molecular weights (10 kDa, 70 kDa, and 500 kDa), and their corresponding hydrogels were dissolved in Milli-Q water and analysed by MALDI-TOF MS to elucidate the polymer composition and confirm repeating monomeric structures. Samples were drop-cast onto a MALDI target plate and dried under ambient conditions. A matrix composed of Sodium Trifluoroacetate and Dihydroxybenzoate in tetrahydrofuran solvent was used. Mass spectra were recorded on a Bruker Autoflex MALDI-TOF instrument operated in linear positive mode across an m/z range of 300–11,000. Data were processed using PolyTools software to identify polymeric peaks and assign repeating-unit masses.

### Thermogravimetric Analysis (TGA) of Hydrogel

Thermogravimetric analysis (TGA) was performed on both freshly formed and lyophilized hydrogels using a NETZSCH STA 449 F3. Samples were heated from 25 °C to 550 °C at a rate of 5 K min□¹ under a nitrogen flow of 50 mL min□¹.

### Confocal Microscopy of Hydrogels

Samples were prepared by combining 5 µL of super folder Green Fluorescent Protein (sfGFP) bacterial culture into dextran, followed by the addition of PEG 400 to spin the mixture into a hydrogel. The resulting hydrogel was gently flattened into a thin sheet using a clean coverslip and then mounted on a glass slide for imaging. Observations were made using 10× and 63× objectives, and data were acquired at 488 nm at the objective lens. Confocal imaging was performed using a Leica TCS SP8 laser scanning confocal microscope (Leica Microsystems, Germany) equipped with an argon laser for 488 nm excitation and a hybrid detector (HyD) for fluorescence detection. The analysis was carried out using ImageJ (Fiji) software.

### Molecular Dynamics Simulations of Coarse-grained PEG-dextran mixtures

We employed implicit solvent Langevin dynamics of the coarse-grained polymer model of PEG and DEX to investigate the effect of varying dextran (DEX) molecular weights on the aqueous two-phase diagram. Both PEG and DEX were modelled as bead-spring polymers (**Figure S2**). PEG was modelled as a 12-bead polymer, whereas dextran was modelled with different chain lengths (7, 14, 28, 40) to examine how the density of the two components in the two phases changes as a function of the chain length. Beads of the two polymers interacted with a purely repulsive Lennard-Jones (LJ) potential (Weeks-Chandler-Andersen potential) as shown in the equation below:

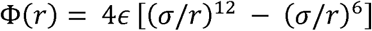

Where Φ(r) is the standard Lennard-Jones potential, r is the distance between the centres of particles *i* and *j*, σ characterizes the distance below which there is strong repulsion and ε denotes the repulsive strength parameter.^27^ We chose PEG–PEG and DEX–DEX, interactions that are identical (pure excluded volume) between the beads of the polymers, while having larger repulsive interactions between PEG-DEX pair of beads to model mutual incompatibility. We used a finitely extensible non-elastic (FENE) potential between adjacent monomer beads with K = 20 ε/σ^2^ and R_0_ = 1.5σ. The table lists the LJ parameters used in the Langevin dynamics simulations.

**Table 1.**
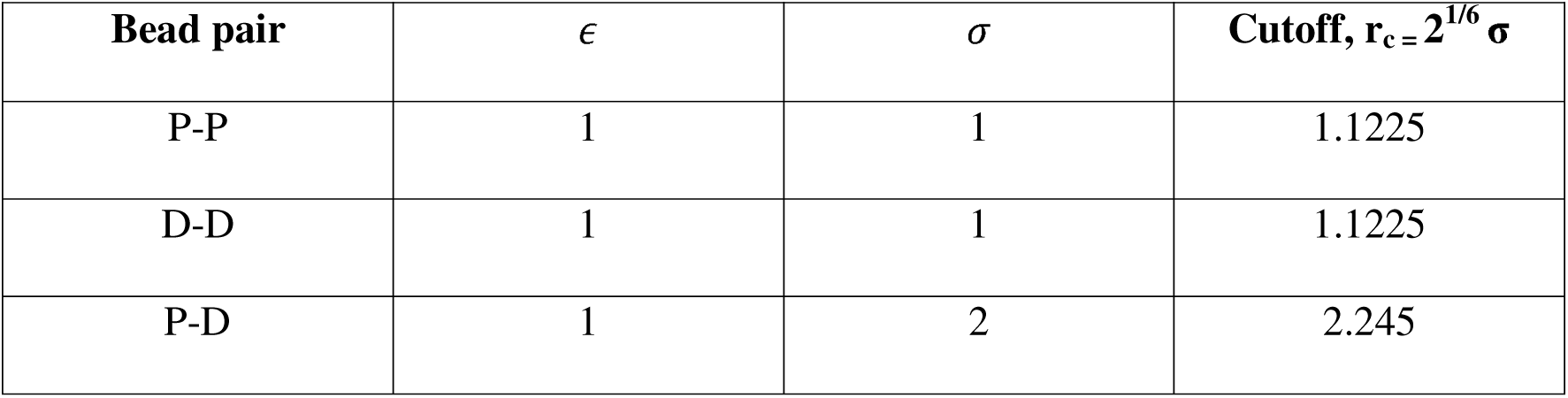
Definition of non-bonded interactions between the two types of beads of the PEG-DEX system.

All the simulations were performed in the NVT ensemble using the LAMMPS (Large-scale Atomic/Molecular Massively Parallel Simulator) simulation package (23 Jun 2022) with reduced Lennard-Jones units where energy, length, mass, and time are nondimensionalized by ε, σ, and m.^27, 28^ The simulation system consisted of a cuboidal box (in slab geometry) with dimensions of 135σ × 40σ × 40σ, containing PEG polymers and DEX polymers. The slab geometry has been well established to study phase separation.^29, 30^ The Langevin equation was integrated forward in time using the Velocity-Verlet algorithm. The time step for integration was chosen to be 0.001τ, where τ is the natural unit of time. The damping parameter was chosen to be 1τ. PEG and DEX particles were initialised randomly in the cuboidal box, and by placing both polymers segregated to accelerate observation of two coexisting phases. We then performed Langevin dynamics simulations. Simulation trajectories were recorded at 10□ time-step intervals, and each simulation was executed in a total of 10 × 10^7^ time steps to reach steady□state coexisting phases, from which density profiles along the x-axis could be extracted. The equilibrated last 5000 trajectory frames (each sampled 10^4^ time steps apart) were used for calculating the equilibrium densities of the PEG-rich and DEX-rich phases. By calculating the density profile from the centre of mass of DEX polymers, the dilute and dense phase number densities for the two types of beads were determined. By carrying out simulations at different input concentrations, tie lines and phase diagrams for different chain lengths of the coarse-grained DEX polymer were constructed.^31^ We used OVITO to visualize the simulation trajectories.^32^

### Antibiotic Release Activity of Hydrogels

A kanamycin stock solution (500 µg mL□¹) was freshly prepared in sterile milli-Q water. From this, 10 µL of the stock was added to 20 µL of a dextran (DEX, 70 kDa) aqueous solution and mixed thoroughly to ensure homogeneity. Subsequently, 20 µL of polyethylene glycol (PEG 400) was added, vortexed resulting in the formation of the hydrogel. For antimicrobial activity testing, *E. coli* DH5α cells were spread uniformly on LB agar plates. The antibiotic-loaded hydrogel was then placed at the centre of the plate, and the plates were incubated at 37 °C for 18–24 h. Following incubation, the zone of inhibition around the hydrogel was observed to assess antibiotic diffusion and antimicrobial activity. A control hydrogel prepared without kanamycin was used for comparison to confirm that inhibition was solely due to antibiotic release from the matrix.

### Preparation of Poly(2-ethl-2-oxazoline) (PEOx)

Hydrogels were synthesized by combining PEOx with DEX 10 kDa. For hydrogel formation, 20 µL of PEOx and 20 µL of 40 wt.% DEX 10 kDa stock solution was mixed in a 0.5 mL tube. The mixture was vortexed thoroughly for 20–30 seconds to form a hydrogel.

### Seed Germination and Growth in PEG/DEX Gel

Green gram (*Vigna radiata*) seeds were embedded within the hydrogels to study their capacity for supporting germination and growth under simple ambient conditions. The gels were prepared as previously described, then transferred into tube caps to serve as confined growth chambers. Individual seeds were gently placed and partially embedded in the centre of each gel to ensure uniform contact with the matrix. The setup was kept at room temperature (∼25 °C), with hydration supplied by spraying water onto the gel surface daily for 15 days. For controls, individual seeds were placed onto tissue paper moistened with water under identical conditions and hydrated at the same spraying frequency.

## RESULTS AND DISCUSSION

Aqueous two-phase system (ATPS) represents an immiscible biphasic aqueous environment typically formed by the combination of two water-soluble polymers mixed above a certain threshold concentration. This results in phase separation into two distinct phases, enriched in one of the polymers. To determine the threshold concentrations for phase separation, we employed the cloud-point titration method to establish the binodal curves between PEG and DEX with molecular weights of 400 and 70 kDa (**Figure 1A**) and 400 and 500 kDa (**Figure S1A**). As expected, mixtures with PEG (10 wt.%) and DEX (20 wt.%) remained homogenous below the binodal curve (**Figure 1B and S1B**). When the composition (PEG–25 wt.% and DEX–20 wt.%) was shifted above the binodal curve, a clear phase-separated ATPS was observed (**Figure 1C and S1C**). Interestingly, when the PEG concentration was increased to 50 wt.%, a cloudy suspension was observable (**Figure 1D and S1D**). Upon closer examination revealed that this suspension was, in fact, a soft, hydrogel that could be handled (**Figures 1E and S1E**). This unexpected finding indicated that the spontaneous formation of a hydrogel could occur without the need for any polymerization crosslinkers.

**Figure 1.**
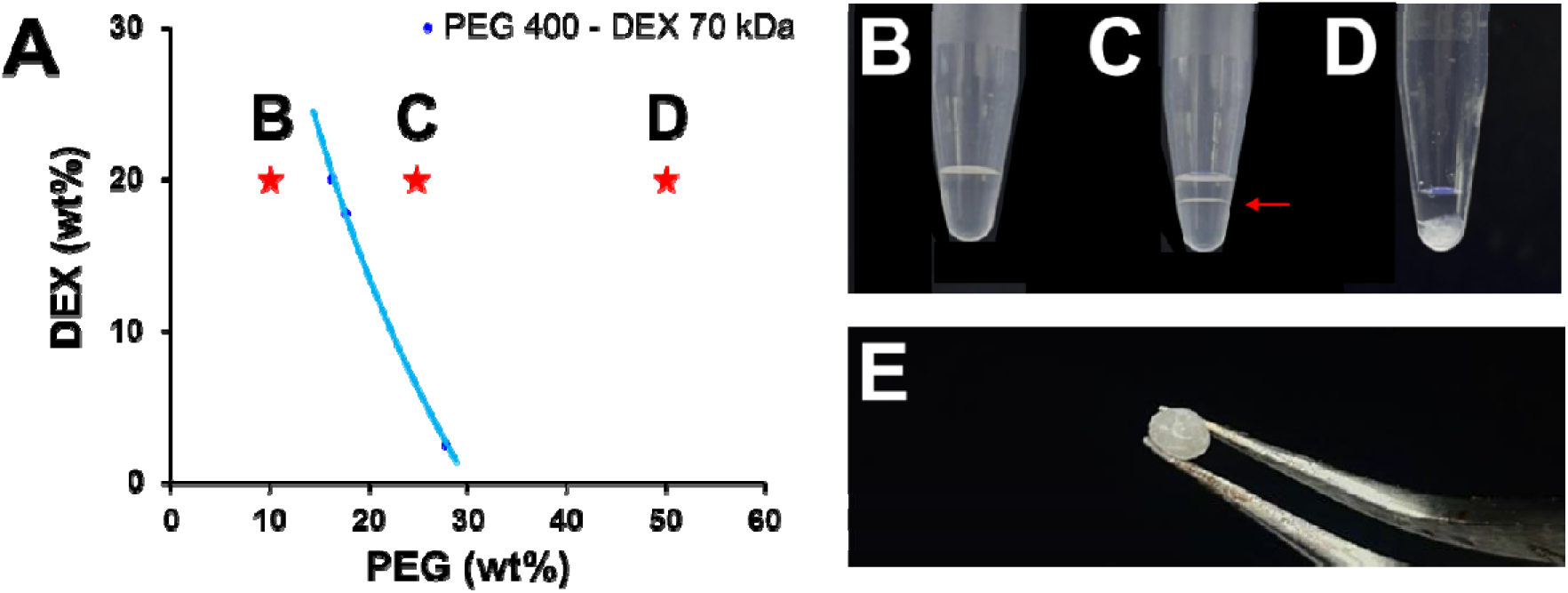
Binodal curve and visual phase separation in PEG 400–DEX 70 kDa systems. **(A)** Binodal curve for PEG 400/DEX 70 kDa system obtained via the cloud-point titration method, showing the boundary between single-phase and two-phase regions, **(B)** Homogeneous phase was observed in the PEG 400 (10%)/DEX 70 kDa (20%) mixture, **(C)** 25% PEG 400/20% DEX 70 kDa two-phase system, with the red arrow indicates the position of the interface, **(D)** 50% PEG 400/20% DEX 70 kDa mixture presenting a cloudy mixture and **(E)** Soft hydrogel is formed when 50% PEG 400 and 20% DEX 70 kDa is mixed and is easy to handle.

To further understand the molecular weight dependence on the formation of these hydrogels, we fixed the molecular weight of PEG and varied the molecular weight of DEX from 10 kDa to 500 kDa. We established the binodal curve for these systems as indicated previously (**Figure 1A and S1A**). The PEG 400/DEX 70 kDa and PEG 400/DEX 500 kDa systems exhibited well-defined binodal boundaries and distinct phase separation. In these systems, the PEG-rich phase formed the upper layer, while the DEX-rich phase constituted the lower layer.^33, 34^ In contrast, for the PEG 400/DEX 10 kDa system, the binodal curve was difficult to delineate, and no discernible phase separation was observed within the investigated concentration range. As DEX 10 kDa and PEG 400 have relatively low molecular weights, which tends to favour mixing over phase separation and push the binodal curve toward higher total concentrations, making phase separation hard to observe.^35–37^ Systems containing higher molecular weight PEG and DEX exhibit more distinct and easily identifiable binodal curves, while systems with lower molecular weight DEX often show weak or suppressed phase separation within typical concentration ranges.

To complement these experimental observations and directly assess the role of polymer chain length on phase behaviour, coarse-grained Langevin dynamics simulations of mutually incompatible polymers were performed (**Figure S2**). We observed that the two-polymer system phase-separated into two coexisting phases: a dense phase rich in dextran (dilute in PEG) and a dilute phase poor in dextran (dense in PEG). Equilibrated snapshots clearly show two coexisting phases separated by an interface (**Figures 2A-D and S3**). **Figure 2E** plots the binodal as a function of different chain length of the dextran polymer. We observed that the two-phase region increases with an increase in dextran chain length. Lower concentration of the two polymers is required to observe phase separation with a higher chain length (molecular weight) dextran. These binodal trends align with the expectation that increasing polymer chain length lowers the critical concentration. We plotted the tie-lines characterizing the densities in the two phases and observed that an increase in chain length leads to more exclusion of PEG from DEX-rich phases and vice-versa. We calculated density profiles along the longer-direction of the slab simulation box to characterize the densities in the two phases. As the chain length of the dextran polymers (shown in blue) increases, more dextran is enriched in the dense phase. Conversely, PEG (shown in red) is increasingly excluded from this dextran-rich phase as the molecular weight of Dextran (N_D_) increases (**Figures 2F and S4**). This behaviour arises because the thermodynamic driving force for demixing grows with polymer chain length, in line with predictions from Flory–Huggin’s theory.^29^ We also observed that the interface sharpness increases upon changing the chain length of dextran. Therefore, our simulations show that to obtain dextran-rich hydrogels, experiments should utilize a high molecular weight dextran. The results from simulations performed at selected volume fractions remain qualitatively comparable to the experimental data, reinforcing that the system undergoes a physical, equilibrium phase separation rather than chemical gelation to stabilize these dense phases.

**Figure 2.**
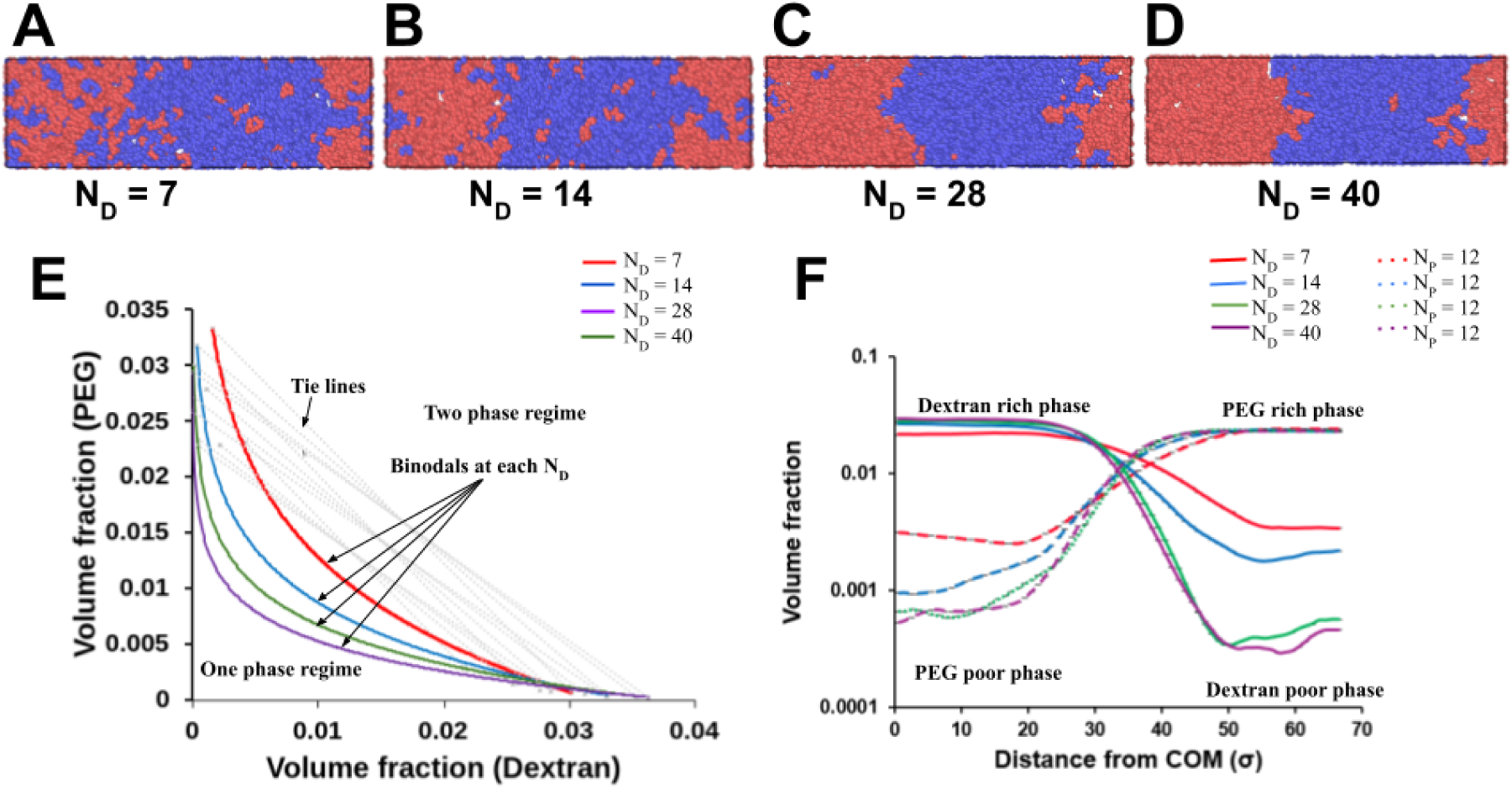
Langevin dynamics simulations illustrating the effect of dextran chain length on PEG (red) – DEX (blue) phase separation. PEG is taken as a 12-mer polymer (N_P_=12), whereas the dextran chain length is varied. **(A-D)** show the representative equilibrium snapshots of slab simulations at Φ_PEG_ = 0.0116 and Φ_DEX_ = 0.0135 for (**A)** N_D_ **=** 7, **(B)** 14, **(C)** 28, and **(D)** 40, respectively. **(E)** Binodal is plotted at different volume fractions (Φ = 0.025, 0.028 and 0.031) with different N_D_ values. As N_D_ increases, binodal shifts downwards showing an increase in the two-phase region. **(F)** VolumeDfraction of PEG and DEX beads as a function of x-distance from centre-of-mass of dextran at Φ = 0.025. It shows that as N_D_ increases, the dextranDrich region in the centre of the slab becomes denser in dextran and more strongly depleted in PEG.

We verified the formation of the hydrogel in these systems at a fixed PEG (50 wt.%) and DEX (20 wt.%) concentrations of varying molecular weights (**Figures 3A-C**). Interestingly, these gels varied in their optical properties (transparency). An increase in the molecular weight of DEX led to a gradual improvement in the optical transparency of the PEG 400–DEX hydrogels. The hydrogel containing DEX 10 kDa appeared opaque (**Figure 3D**). The DEX 70 kDa hydrogel showed translucency (**Figure 3E**). In contrast, the DEX 500 kDa hydrogel was highly transparent, allowing clear visibility of the underlying text (**Figure 3F**). After being left outside for seven days, the gels showed minimal signs of progressive dehydration with a slight decrease in transparency (**Figures 3G-I**).

**Figure 3.**
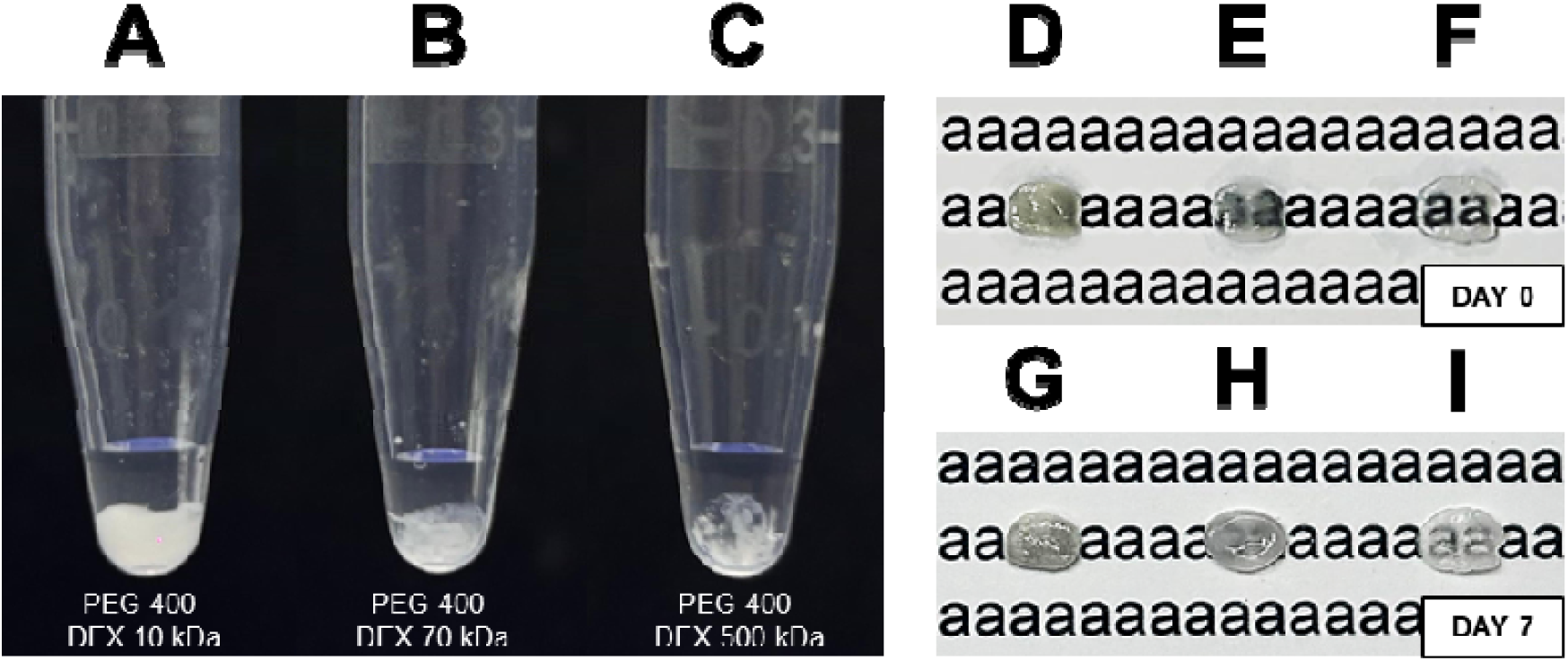
Optical clarity assessment of PEG 400/DEX hydrogels. **(A–C)** Formation of PEG 400/DEX hydrogels of different molecular weights (10 kDa, 70 kDa, and 500 kDa) observed immediately after mixing, **(D–F)** Optical transparency assessment of freshly prepared hydrogels (Day 0) placed over printed text. **(D)** Hydrogel A (DEX 10 kDa) appeared opaque, **(E)** Hydrogel B (DEX 70 kDa) showed partial translucency, and **(F)** Hydrogel C (DEX 500 kDa) exhibited high transparency and **(G–I)** Corresponding hydrogels at Day 7, exhibiting decreased transparency likely due to gradual dehydration.

To determine the compositional nature of the gel—whether it was enriched in PEG or DEX—we incorporated a dye during the gelation process (**Figures 4AI, BI, and CI**). The dye incorporated hydrogel when placed in a PEG 400 solution, resulted in no dissolution of the hydrogel (**Figures 4AII, BII, and CII**). In contrast, when immersed in DEX solution of the same molecular weight, the hydrogel readily dissolved (**Figures 4AIII, BIII, and CIII**). These suggest that the hydrogel was predominantly DEX-rich. We additionally examined the supernatant after the formation of the hydrogel and the hydrogel dissolved in water using a colorimetric assay specific for PEG (**Figures S5**). In this method, the solution is mixed with ammonium ferrothiocyanate and chloroform, producing a red colour in the chloroform layer when PEG is present. A strong red colouration of the supernatant validates PEG enrichment (**Figure S5D**), while the gel showed minimal PEG (weak red coloration), indicating it was mainly dextran-rich (**Figure S5E**). The ratio of PEG concentration in the supernatant to that in the gel was determined to be ∼8:1 (**Figure S5F**).

**Figure 4.**
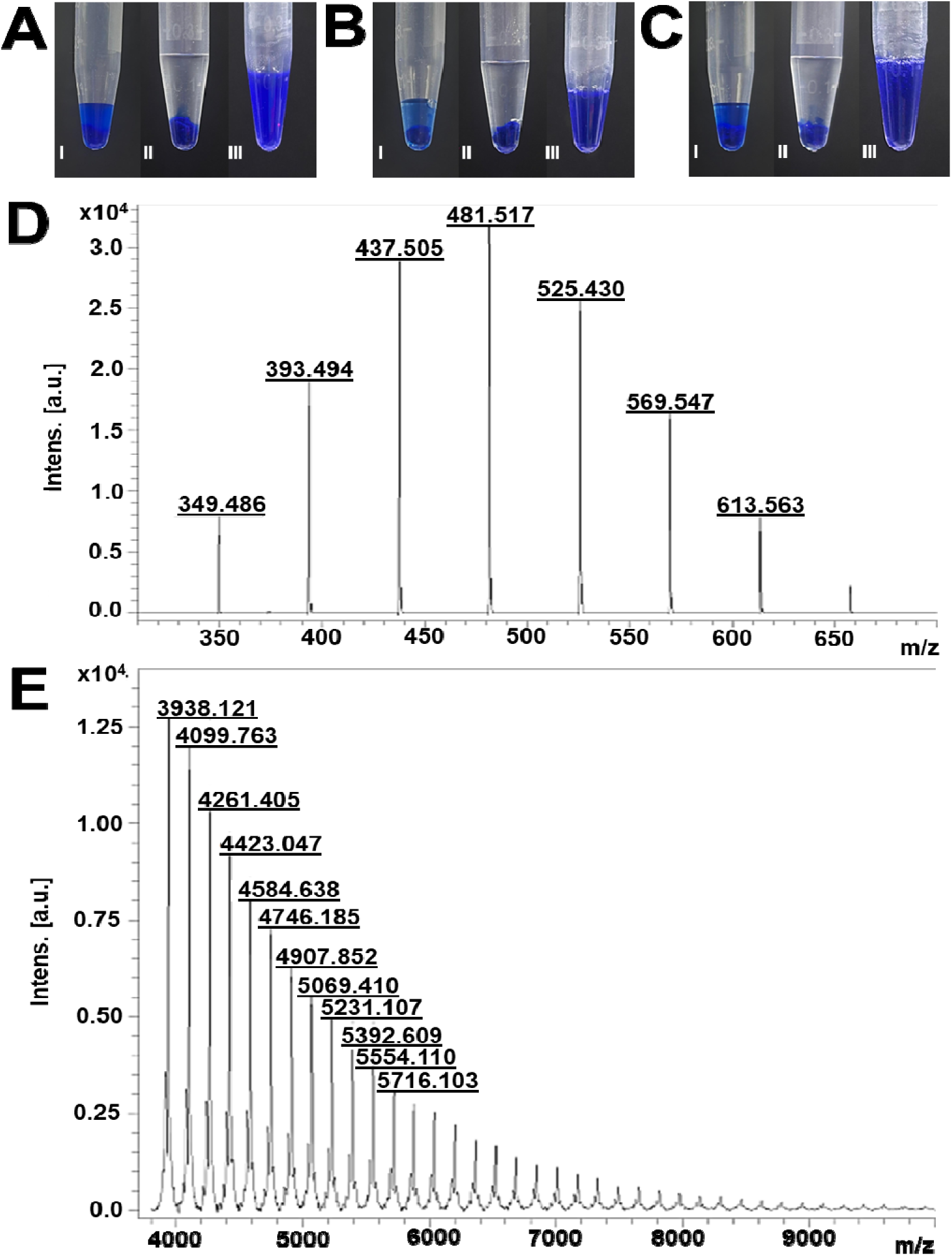
Solubility behaviour and compositional analysis of PEG 400–DEX systems of varying molecular weights. **(A–C)** Representative images showing hydrogel formation using PEG 400 with DEX of increasing molecular weights: **(A)** 10 kDa, **(B)** 70 kDa, and **(C)** 500 kDa, **(I)** Coomassie Brilliant Blue dye incorporated within PEG and DEX mixtures, **(II)** Blue dyed hydrogel immersed in excess PEG exhibiting no dissolution, and **(III)** Blue dyed hydrogel immersed in excess of DEX showing complete dissolution, MALDI-TOF mass spectra of **(D)** Supernatant of PEG 400/DEX 10 kDa hydrogel, **(E)** Hydrogel fabricated by PEG 400/DEX 10 kDa dissolved in water. The supernatant spectra display characteristic repeating units at ∼44 Da intervals, corresponding to ethylene oxide monomers of PEG, which confirms PEG enrichment in the upper phase. Conversely, the hydrogel spectrum shows ∼162 Da intervals associated with D-glucose residues, indicating dextran predominance in the condensed phase.

These findings were corroborated by MALDI-TOF mass spectrometry (**Figures 4D-E and S6A-J**). The spectra of pure PEG 400 (**Figure S6A**), along with its characteristic ethylene oxide monomeric repeating units at ∼44 Da (**Figure S6B**). The supernatant spectra obtained from all PEG/DEX systems (**Figures 4D, S6C, and S6E**) showed peak patterns matching those of PEG 400, and their corresponding repeating-unit analyses (**Figures S6D, S6F, and S6I**) further confirmed PEG enrichment in the upper phase. Conversely, the hydrogel (formed with PEG 400/DEX 10 kDa) and dissolved in water revealed peaks corresponding to dextran 10 kDa (**Figures 4E and 6G, S6H and S6J**) corresponding to monomeric D-glucose (∼162 Da) repeating units of dextran. These findings corroborate that the hydrogels are primarily DEX-rich, while the supernatant phase is PEG-rich.

The dissolution behaviour of these gels was evaluated by immersing the dyed hydrogels in water and monitoring their disintegration over time (**Figure 5**). The dissolution kinetics exhibited a clear dependence on dextran molecular weight. Hydrogels composed of PEG 400/DEX 10 kDa (**Figure 5A**) and PEG 400/DEX 70 kDa (**Figure 5B**) completely dissolved within ∼25 minutes, whereas hydrogels formed from PEG 400/DEX 500 kDa (**Figure 5C**) required approximately 50 minutes for complete dissolution. Quantitative analysis of the reduction in hydrogel diameter over time confirmed this trend, with higher-molecular-weight dextran imparting greater resistance to dissolution (**Figure 5D**). The gradual rise in dissolution time with increasing dextran molecular weight is likely due to the establishment of a denser and more entangled polymeric network at elevated molecular weights.^29, 38^

**Figure 5.**
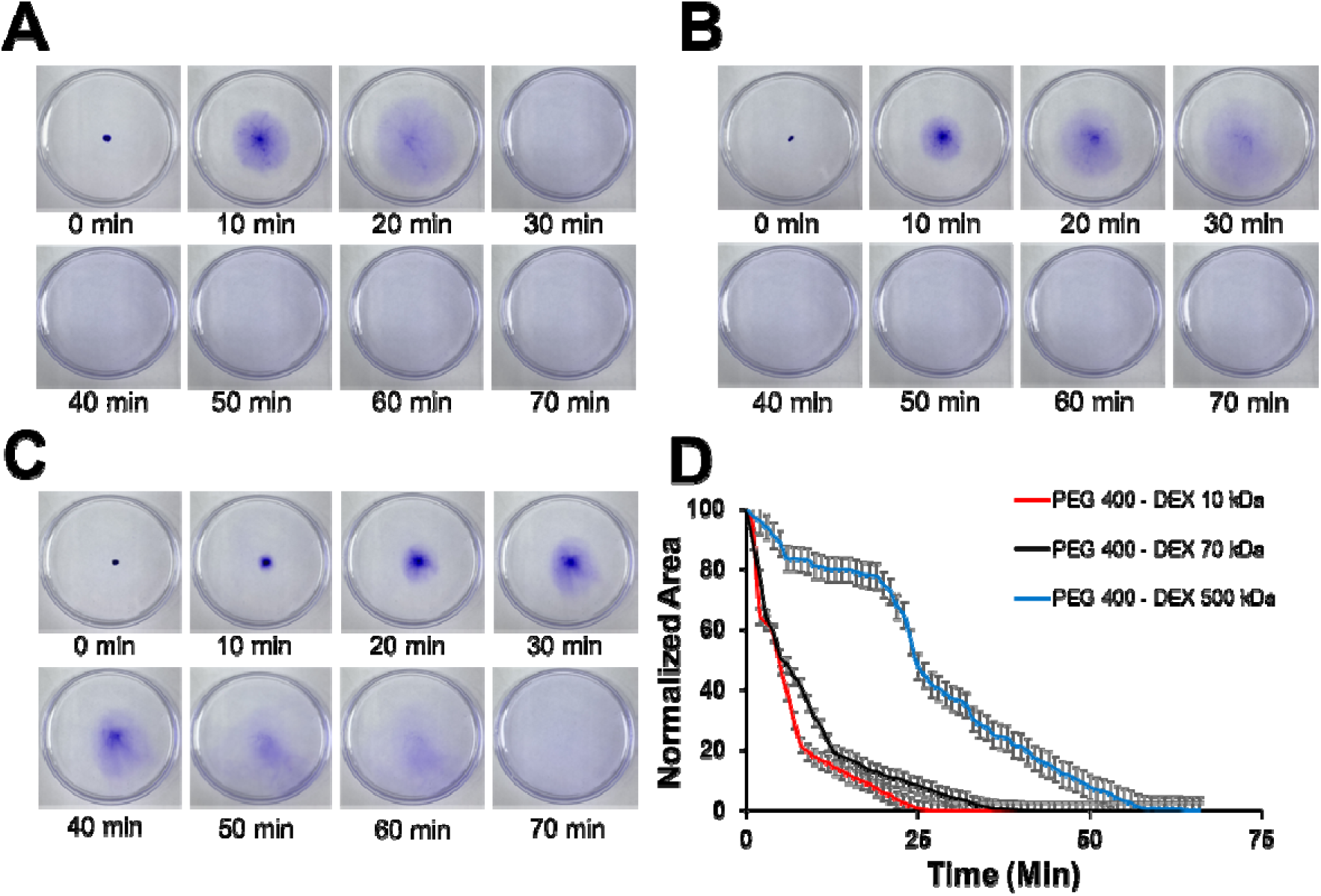
Dissolution kinetics of PEG–DEX gels visualized using Coomassie Brilliant Blue dye. Hydrogels prepared using different DEX molecular weights were immersed in 25 mL of Milli-Q water. Time-lapse images show dissolution and dye release for **(A)** PEG 400/DEX 10 kDa, **(B)** PEG 400/DEX 70 kDa, and **(C)** PEG 400/DEX 500 kDa hydrogels. **(D)** Corresponding dissolution profiles were obtained by quantifying the decrease in hydrogel size over time.

A schematic illustration of the PEG–DEX hydrogel (**Figure 6A**), depicting the entrapment of PEG and water within the dextran-rich matrix. To further examine solvent distribution, volume fraction profiles of the solvent from the simulations were derived to study solvent entrapment in the dense phase (**Figure 6B**). Solvent fraction was calculated as Φ_SO1vent_ == 1 - Φ_DEX_ - Φ_PEG_ at all the N_D_ values. The solvent volume fraction in the DEX-rich phase ranges from 0.975 to 0.970. The interface region shows an increased solvent presence due to low density of the either of the two polymers, owing to their mutual incompatibility. **Figure 6B** shows the presence of solvent in the dextran rich phase, which is further validated with Thermogravimetric analysis (TGA) analysis. TGA of lyophilized and non-lyophilized PEG 400/dextran hydrogels (**Figures 6C-E and S7**) provided clear evidence of solvent entrapment within the polymer matrix. The non-lyophilized samples showed an initial mass loss below ∼100°C, which can be attributed to the removal of trapped water.^39^ In comparison, the lyophilized samples showed almost no mass loss in this region, confirming that free water was effectively removed during drying. A gradual mass loss extending up to ∼200°C was also seen in the non-lyophilized samples, suggesting the presence of retained PEG 400, as PEG starts to degrade around 200–250°C while dextran begins to decompose above ∼280°C.^40, 41^ These results indicate that both water and PEG were partially retained within the gel. Supporting this, FTIR spectra showed a distinct PEG ether (C–O–C) stretching peak near 1098 cm□¹, confirming that PEG was incorporated into the dextran matrix (**Figure S8**).

**Figure 6.**
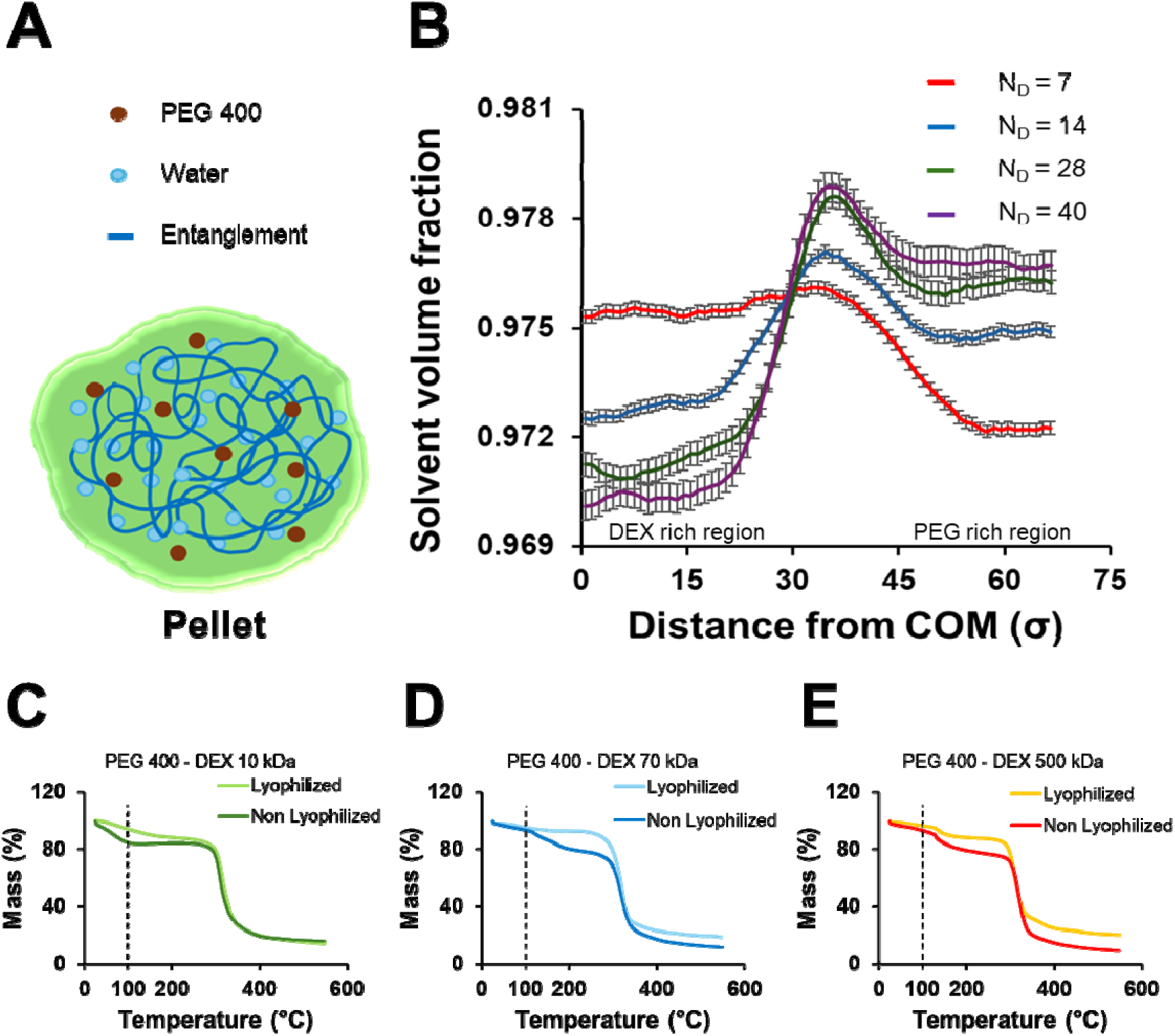
Solvent distribution and thermal characteristics of PEG–DEX hydrogels. **(A)** Schematic representation of the PEG–DEX hydrogel illustrating the entrapment of PEG and solvent (water) within the polymer-rich condensed phase. **(B)** Solvent volume fraction profiles obtained from Langevin dynamics simulations, plotted as a function of distance from the centre of mass along the x-axis for varying dextran chain lengths. The profiles indicate that solvent remains present within both the DEX-rich and PEG-rich regions, confirming heterogeneous solvent distribution throughout the hydrogel. **(C–E)** Thermogravimetric analysis (TGA) of lyophilized and non-lyophilized hydrogels shows an initial mass loss at ∼100 °C due to trapped water, whereas lyophilized hydrogels show minimal loss.

Together, these results confirm that both water and PEG, remained trapped within the dextran-rich gel due to close polymer–polymer interactions that reduced complete separation. Together, these results confirm that both water and PEG, remained trapped within the dextran-rich gel due to close polymer–polymer interactions that reduced complete separation.

Building on our earlier fundamental characterization of the hydrogel system, we next sought to demonstrate its functional performance. We successfully entrapped super-folder green fluorescent expressing bacteria within the hydrogel matrix (**Figure 7A-B**). The cells retained their cylindrical morphology, highlighting minimal cell stress while remaining physically confined within the hydrogel. In a separate experiment, we loaded kanamycin into the PEG 400/DEX 70 kDa hydrogel and assessed their antibiotic release capability using a zone of inhibition assay (**Figures 7C-E**). Hydrogel containing the antibiotic produced a distinct and clear inhibition zone around the dissolved gel, indicating effective antibiotic release from the matrix (**Figure 7E**). In contrast, the control hydrogel lacking antibiotic showed no inhibition zone and exhibited normal bacterial growth around the hydrogel area (**Figure 7D**). Finally, we entrapped plant seeds within these hydrogels (**Figure 7G**). These hydrogels could effectively support seed germination and growth under ambient conditions. Together, these results confirm that the hydrogel functions as a reliable entrapment vehicle for small molecules, living cells and plant seeds. To expand the potential of this technique, we explore PEOx to phase separate with DEX. This combination also resulted in the formation of hydrogels (**Figure S9**).

**Figure 7.**
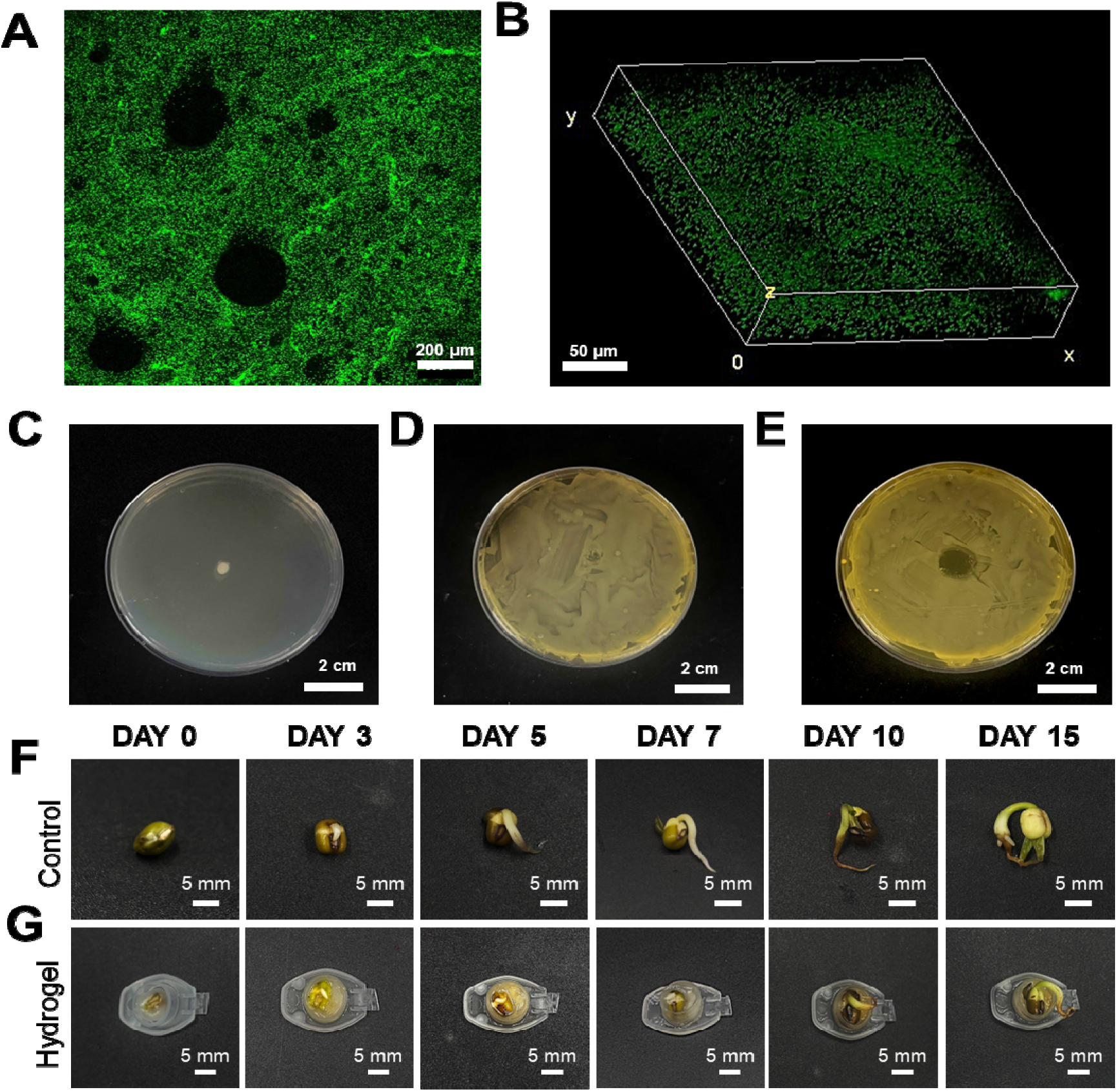
Hydrogels are employed to encapsulate cells, load antibiotics, and as a seed growth support. **(A)** Confocal laser scanning microscopy (z-stack) image showing uniform distribution of sfGFP-expressing bacteria (green fluorescence) entrapped within the hydrogel, **(B)** Three-dimensional reconstruction of the z-stack confirms homogeneous confinement of bacteria, **(C)** Hydrogels placed on an agar plate at t = 0, **(D)** Hydrogel without antibiotic showing no zone of inhibition, **(E)** Hydrogel with antibiotic showing a distinct clear zone of inhibition, **(F)** Control seeds grown on moistened tissue paper and **(G)** Seeds embedded in PEG 400/DEX 500 kDa hydrogels showing successful germination and growth under ambient conditions for 15 days.

## CONCLUSIONS

To the best of our knowledge, this study reports a spontaneous, polymerization-free strategy for the fabrication of hydrogels through phase separation of polymers. We highlight the potential of this technique by using two GRAS polymers poly (ethylene glycol) (PEG) and dextran (DEX). We demonstrate spontaneous gel formation (3 s) through simple mixing of the polymers significantly away from the binodal curve in the two-phase region. We conducted comprehensive characterization of the hydrogels through FTIR, MALDI-TOF and colorimetric assays which indicated DEX rich hydrogels. Thermogravimetric analysis revealed the presence of water within the hydrogel network. We also demonstrate that the optical transparency of these hydrogels can be fine-tuned based on the molecular weight of dextran. The hydrogels formed with higher molecular weight DEX also demonstrated slower dissolution rates, likely due to greater polymer chain entanglement and reduced mobility. Coarse-grained molecular dynamics simulations provided further insight into the phase separation mechanism, revealing molecular-weight-dependent segregation and the formation of coexisting PEG- and DEX-rich domains that underpin the observed gelation behaviour. Finally, we demonstrate uniform entrapment of green fluorescent protein-expressing bacteria within the hydrogel matrix. Independently, we illustrate sustained antibiotic release through these hydrogels against *E. coli*. Furthermore, seed encapsulation experiments demonstrated successful germination and growth within the hydrogels. Collectively, these results establish PEG–DEX hydrogels as robust, biocompatible carriers capable of encapsulating and providing controlled release of bioagents, offering a versatile and scalable platform for next-generation biomaterials in drug delivery, tissue engineering, and bioactive encapsulation systems.

## AUTHOR CONTRIBUTIONS

The original idea, research concept, and experimental design were developed by K.P. N.P. carried out experiments and data analysis. V.S. carried out molecular simulations and their analysis, and G.C. supervised the molecular simulations and their analysis. A.K. provided feedback and helped shape the analysis and the manuscript. K.P. supervised the overall research and provided critical feedback during manuscript writing. K.P. is thankful to the Department of Biotechnology BT/PR45697/BCE/8/1797/2023 for the project funding.

## ACKNOWLEDGEMENTS

The authors would like to sincerely thank IIT Gandhinagar for providing the facilities and resources necessary to carry out this study. We thank CIF (Central Instrumentation Facility) and CRTDH (Common Resource and Technology Development Hub) for the instrumentation facilities. ChatGPT and Perplexity AI were used for rephrasing the sentences in the manuscript.

