## Supplementary Information for "Spontaneous Phase Separation Enables Rapid, Polymerization-Free Fabrication of Dissolvable Hydrogels"

Karthik Pushpavanam, Ph.D.

Department of Chemical Engineering

Indian Institute of Technology Gandhinagar

Gujarat, India, 382055, India

Gaurav Chauhan, Ph.D.

Department of Chemical Engineering

Indian Institute of Technology Indore

Madhya Pradesh, India, 453552, India

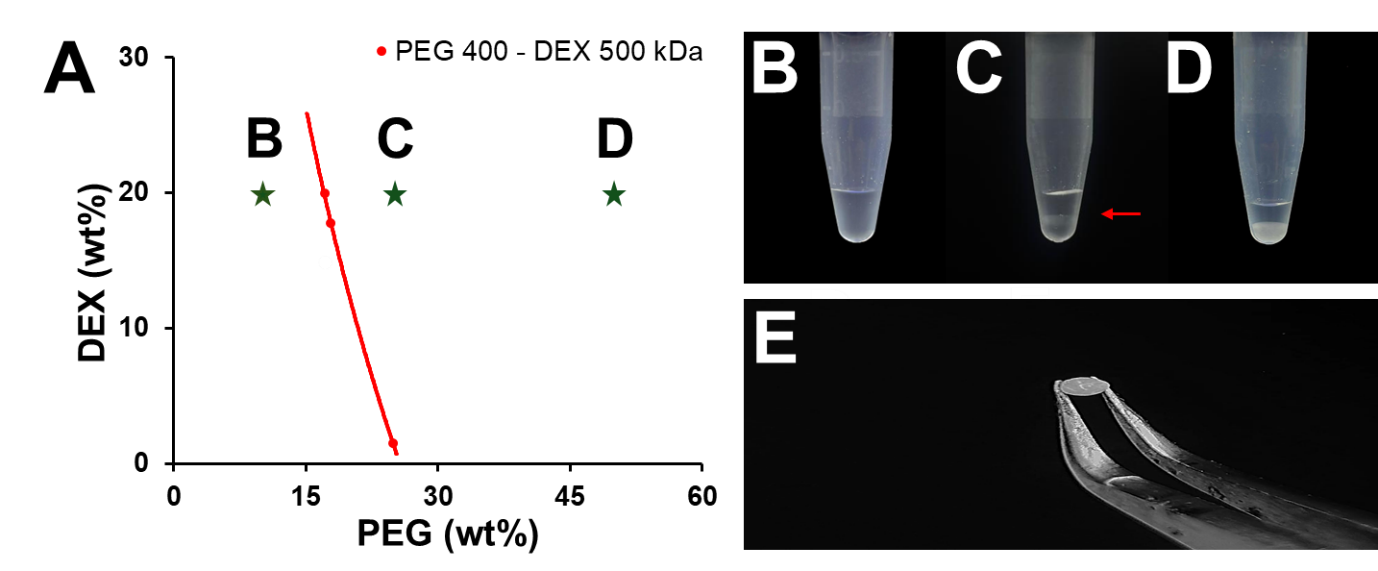

**Figure S1. Binodal curve and visual phase separation in PEG 400–DEX 500 kDa systems. (A)** Binodal curve for PEG 400/DEX 500 kDa system obtained via the cloud-point titration method, showing the boundary between single-phase and two-phase regions, **(B)** Homogeneous phase was observed in the PEG 400 (10%) /DEX 500 kDa (20%) mixture, **(C)** 25% PEG 400/20% DEX 500 kDa two-phase system, with the red arrow indicates the position of the interface, **(D)** 50% PEG 400/20% DEX 500 kDa mixture presenting a cloudy mixture **(E)** Soft hydrogel is formed when 50% PEG 400 and 20% DEX 500 kDa is mixed and is easy to handle.

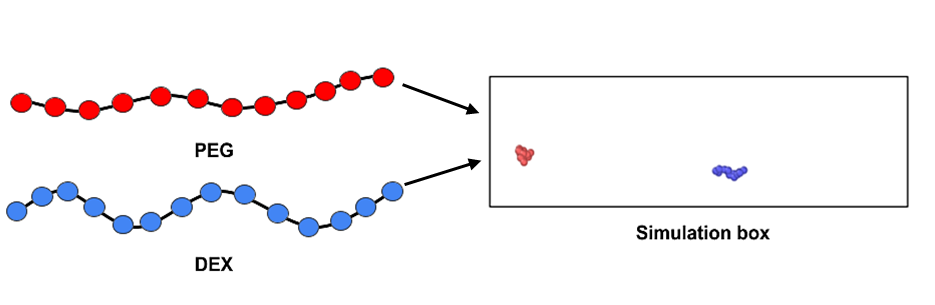

**Figure S2. Coarse-grained model of PEG and DEX particles.** Schematic representation of the coarse-grained model and the simulation setup of the PEG/DEX system. PEG and DEX are considered beads on chains joined through bonds. Copies of these polymers are then introduced into the simulation box.

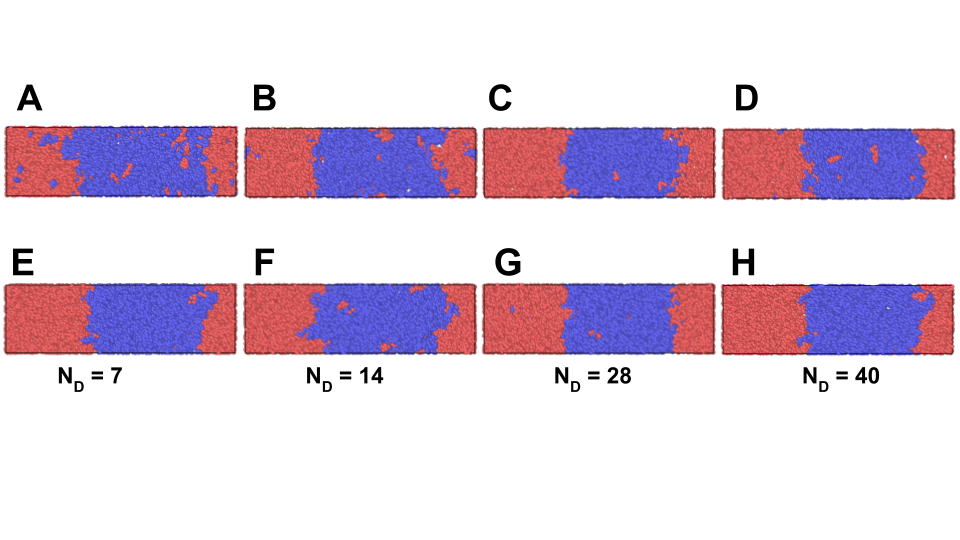
**Figure S3. Illustrative images of the simulated system and the production of ATPS.** Assembly configurations in PEG-DEX systems at different volume fractions with varied dextran chain length, N_D_ = 7 **(A & E)**, 14 **(B & F)**, 28 **(C & G)**, 40 **(D & H)**, and PEG (N_P_ = 12). Here, A-D is at volume fraction 0.028, and E-F is at volume fraction 0.031. Dextran packing becomes more compact in the dense phase as the volume percentage (top to bottom) and chain length (left to right) increases, while more PEG is excluded and embedded in the dilute phase.

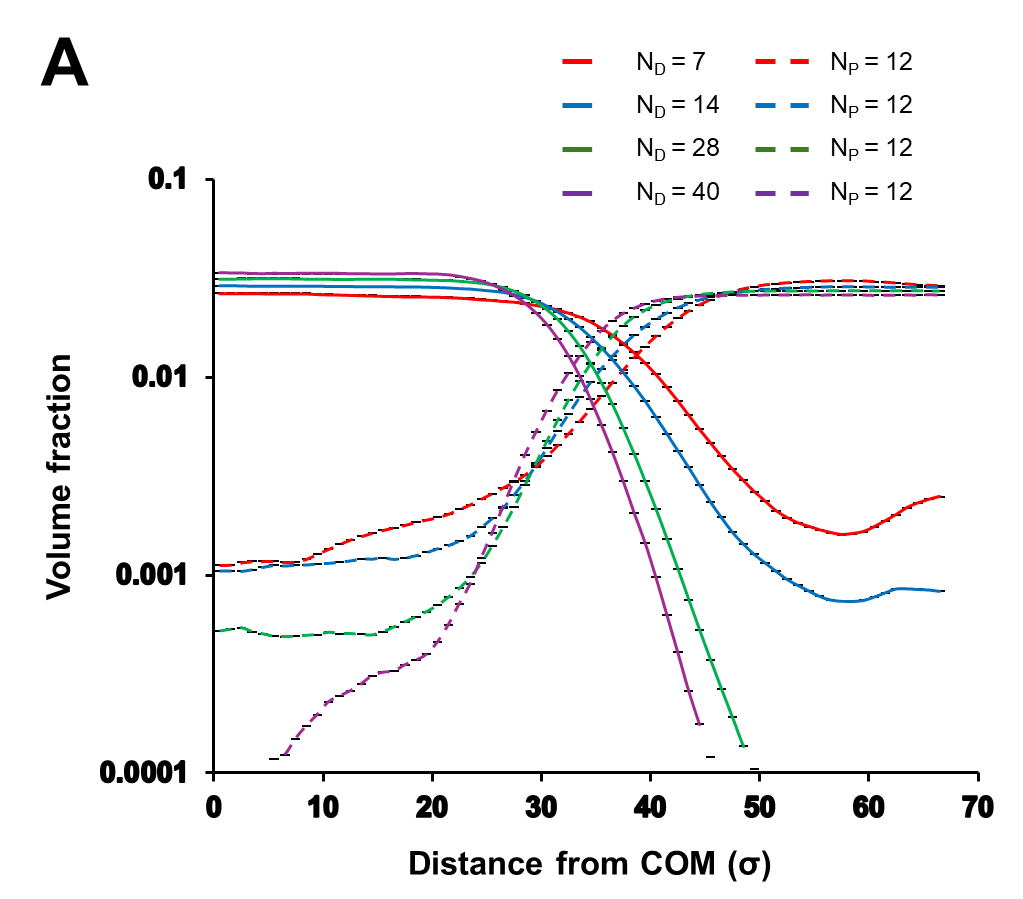

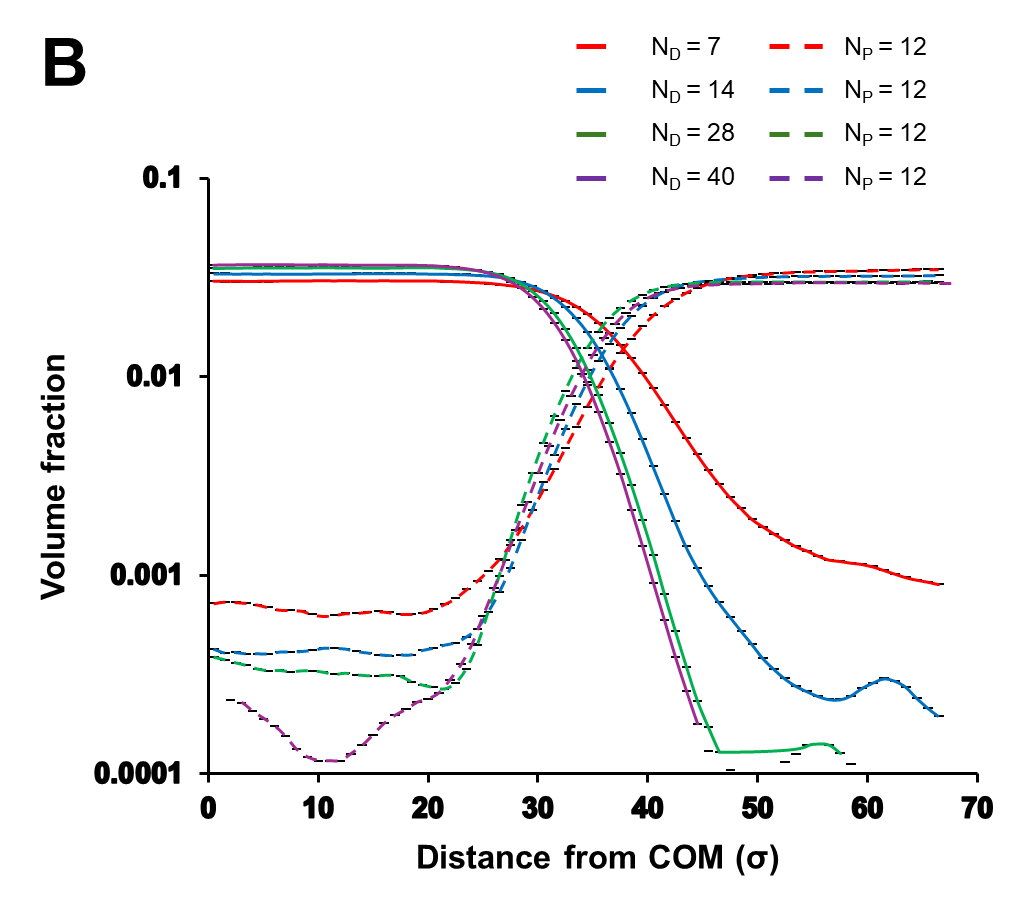

**Figure S4. Volume fraction profiles of PEG and DEX at varying system concentrations.** Volume fraction vs distance plots are shown for DEX (solid lines) and PEG (dashed lines) at different dextran chain lengths (N_D_ = 7, 14, 28, and 40) with a fixed PEG chain length (N_P_ = 12), plotted as a function of the distance from the DEX centre of mass (COM). The graph corresponds to simulations at an overall polymer volume fraction **(A)** ϕₜₒₜ = 0.025, and **(B)** to ϕₜₒₜ = 0.031. The profiles illustrate that longer dextran chains promote enhanced condensate formation and a sharper interface at higher overall concentrations.

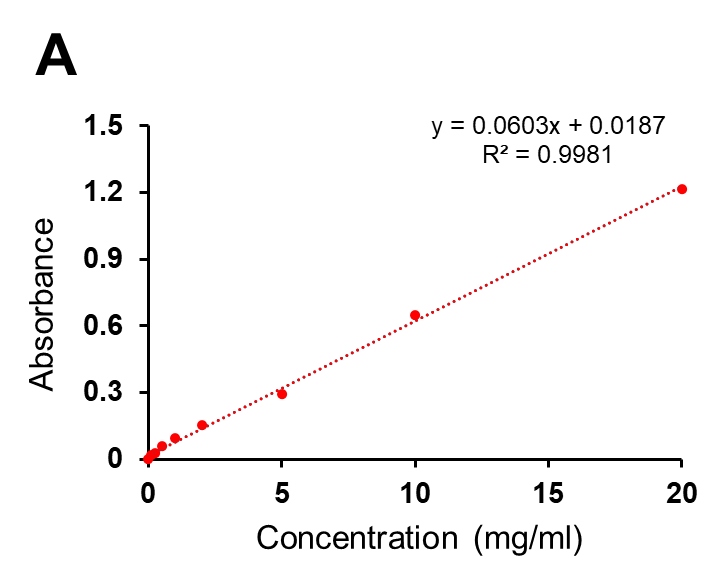

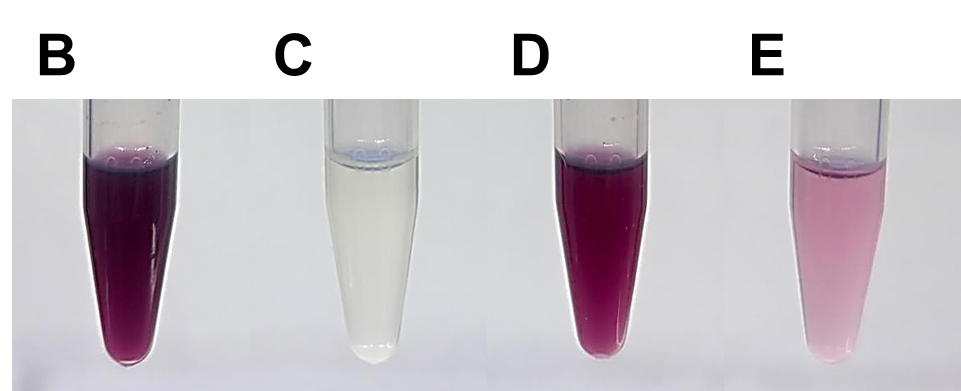

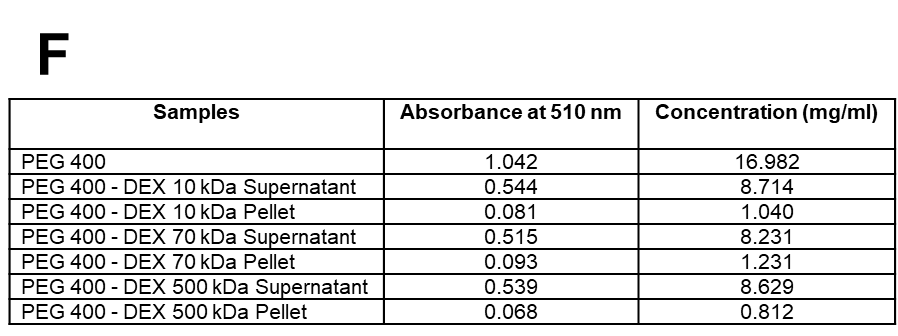

**Figure S5. Colorimetric assay for PEG quantification. (A)** Calibration curve generated using known concentrations of PEG 400, measured at 510 nm with the ammonium ferrothiocyanate–chloroform colorimetric method. **(B–E)** Representative images of the chloroform (lower) phase after extraction, **(B**) Pure PEG 400, **(C)** Dextran 10 kDa, **(D)** Supernatant obtained after hydrogel formation, **(E)** Hydrogel dissolved in Milli-Q water. **(F)** PEG quantification shows an ~8:1 supernatant-to-hydrogel ratio, demonstrating PEG enrichment in the supernatant.

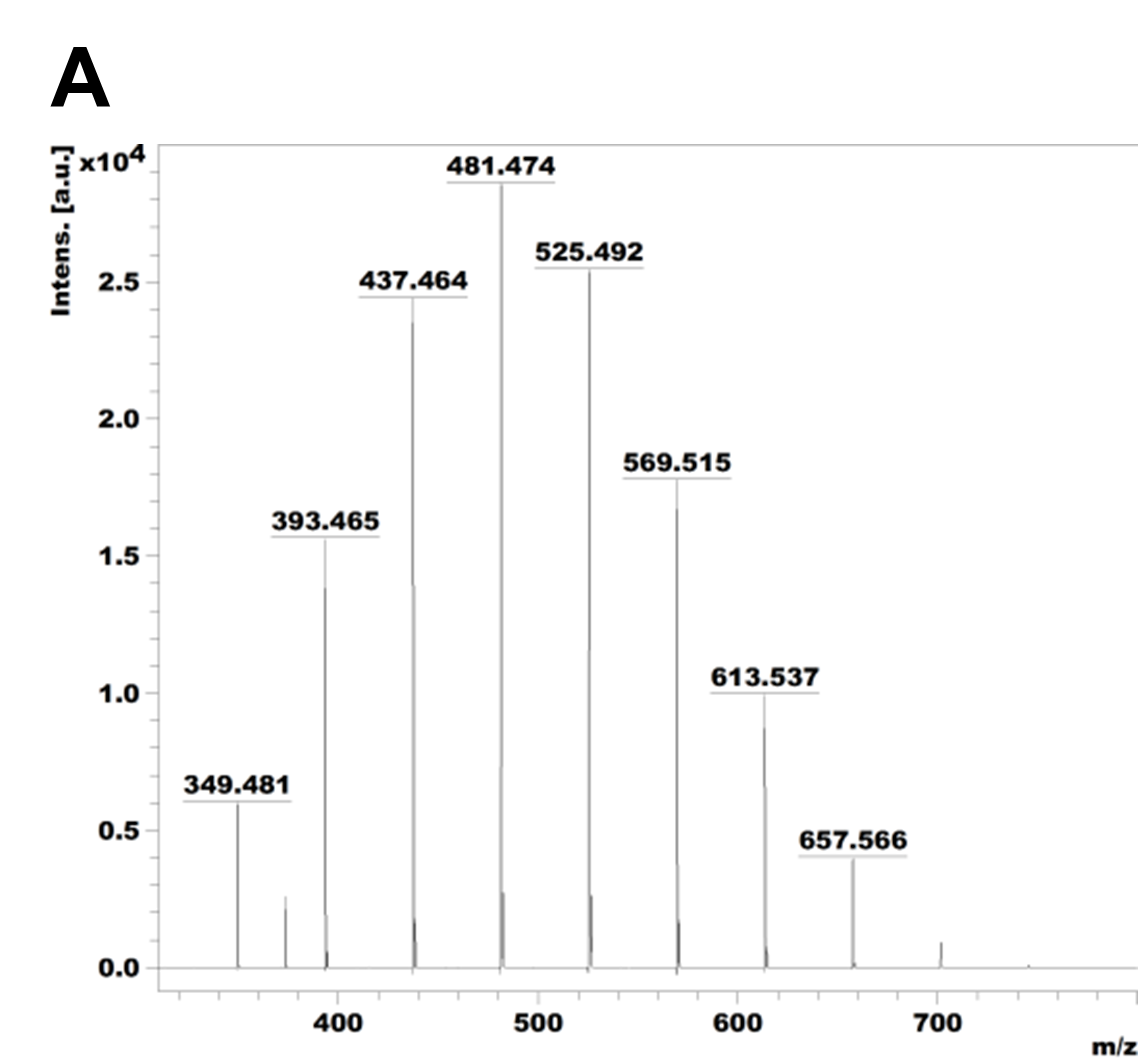

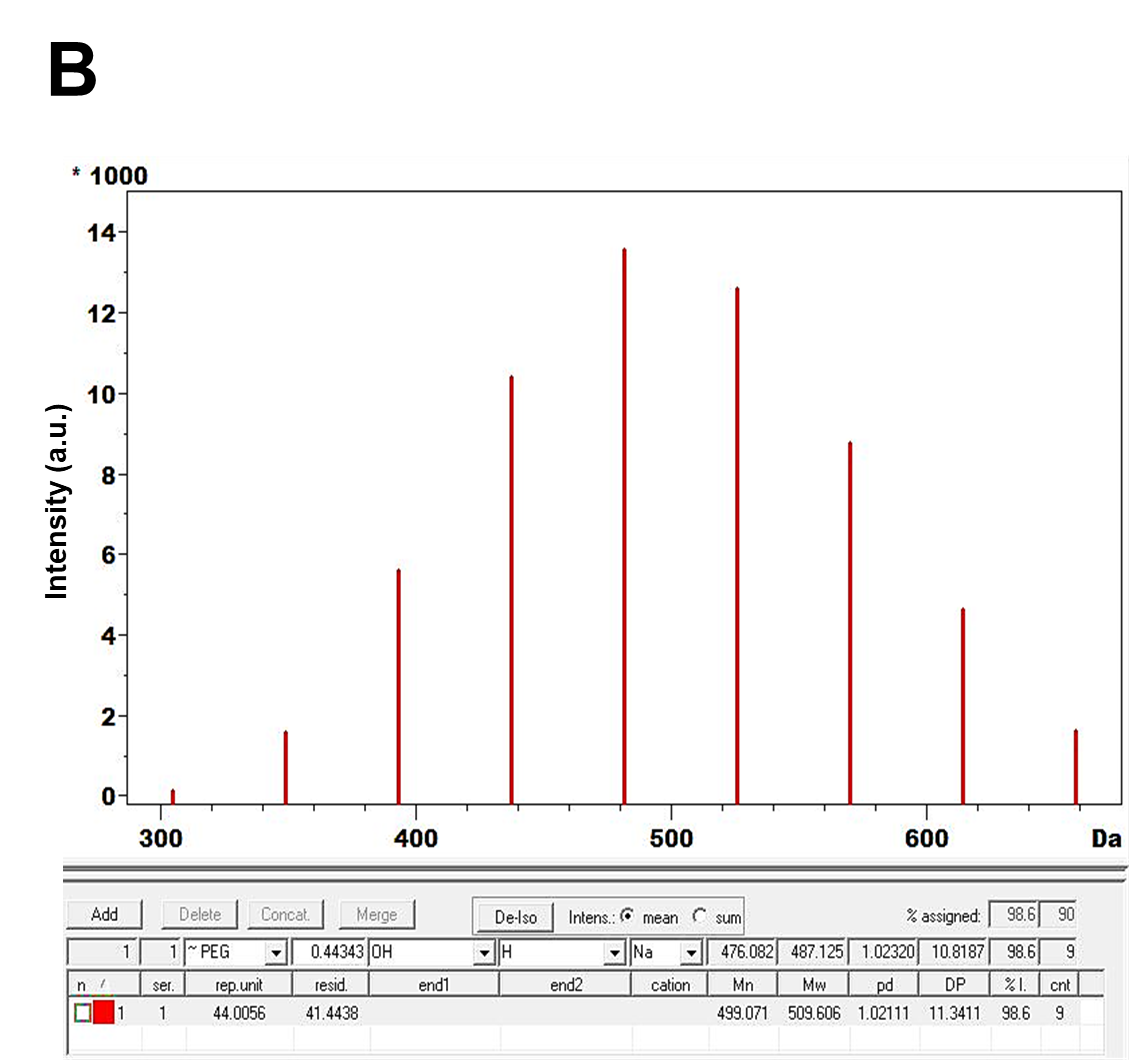

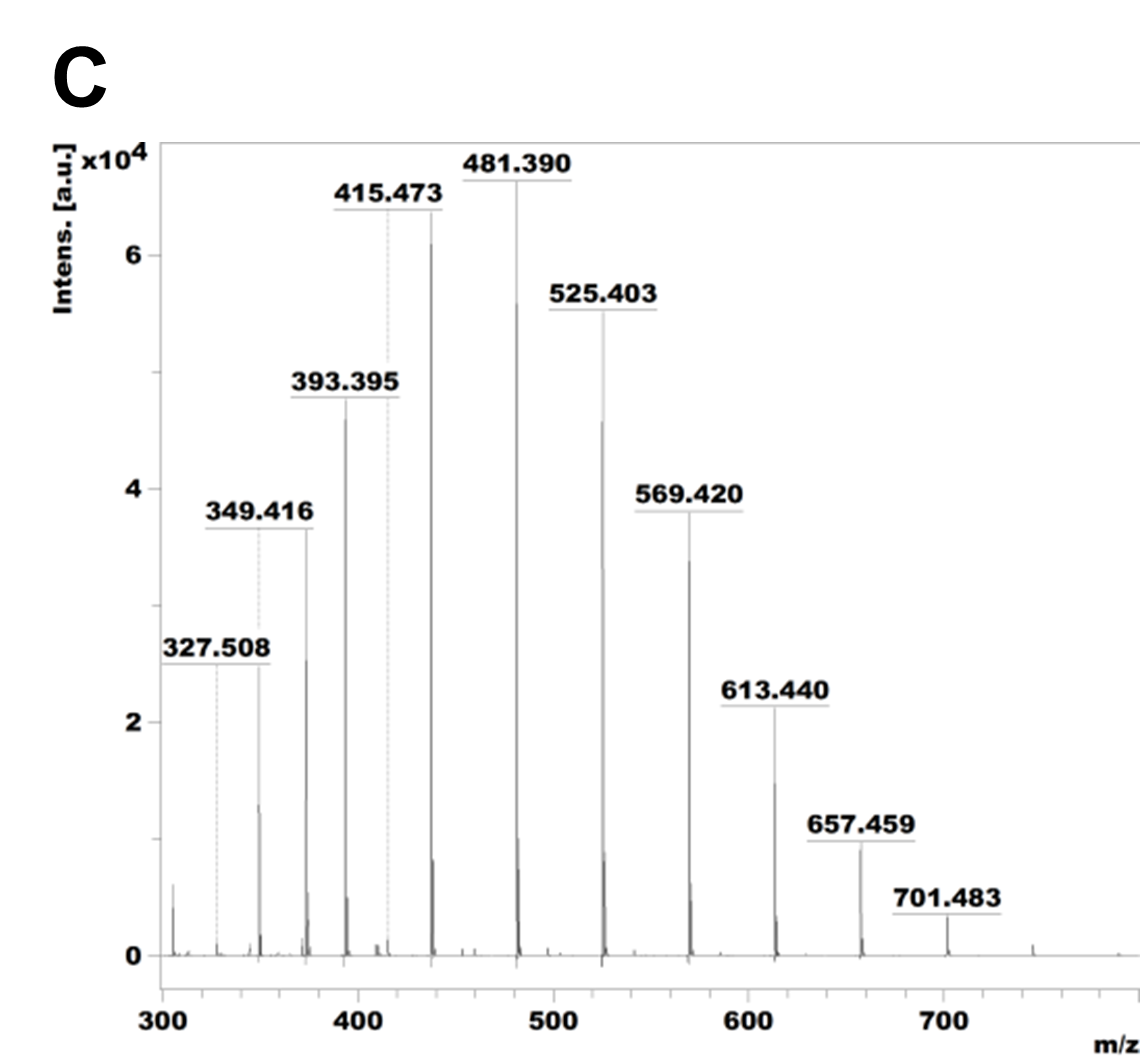

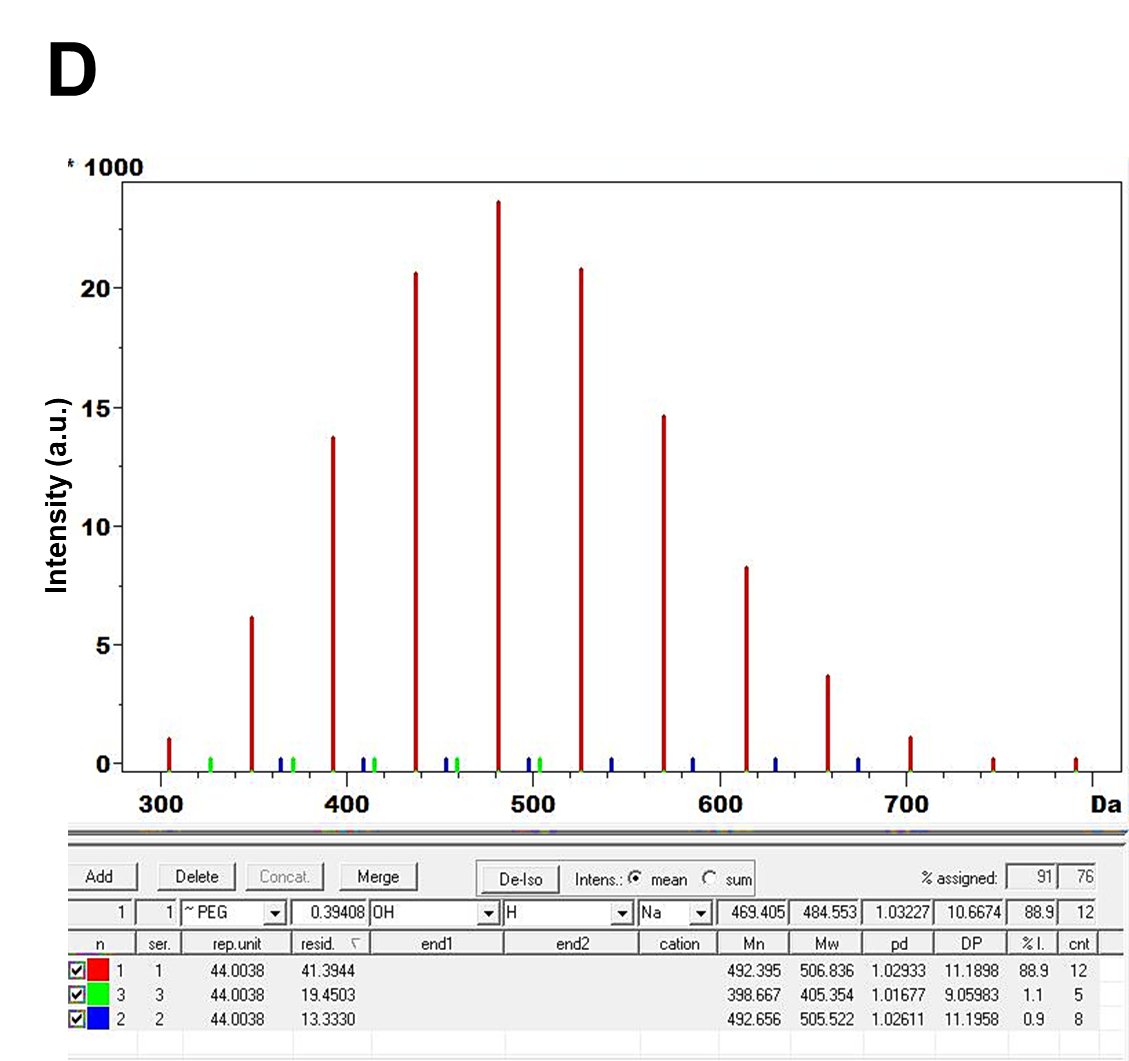

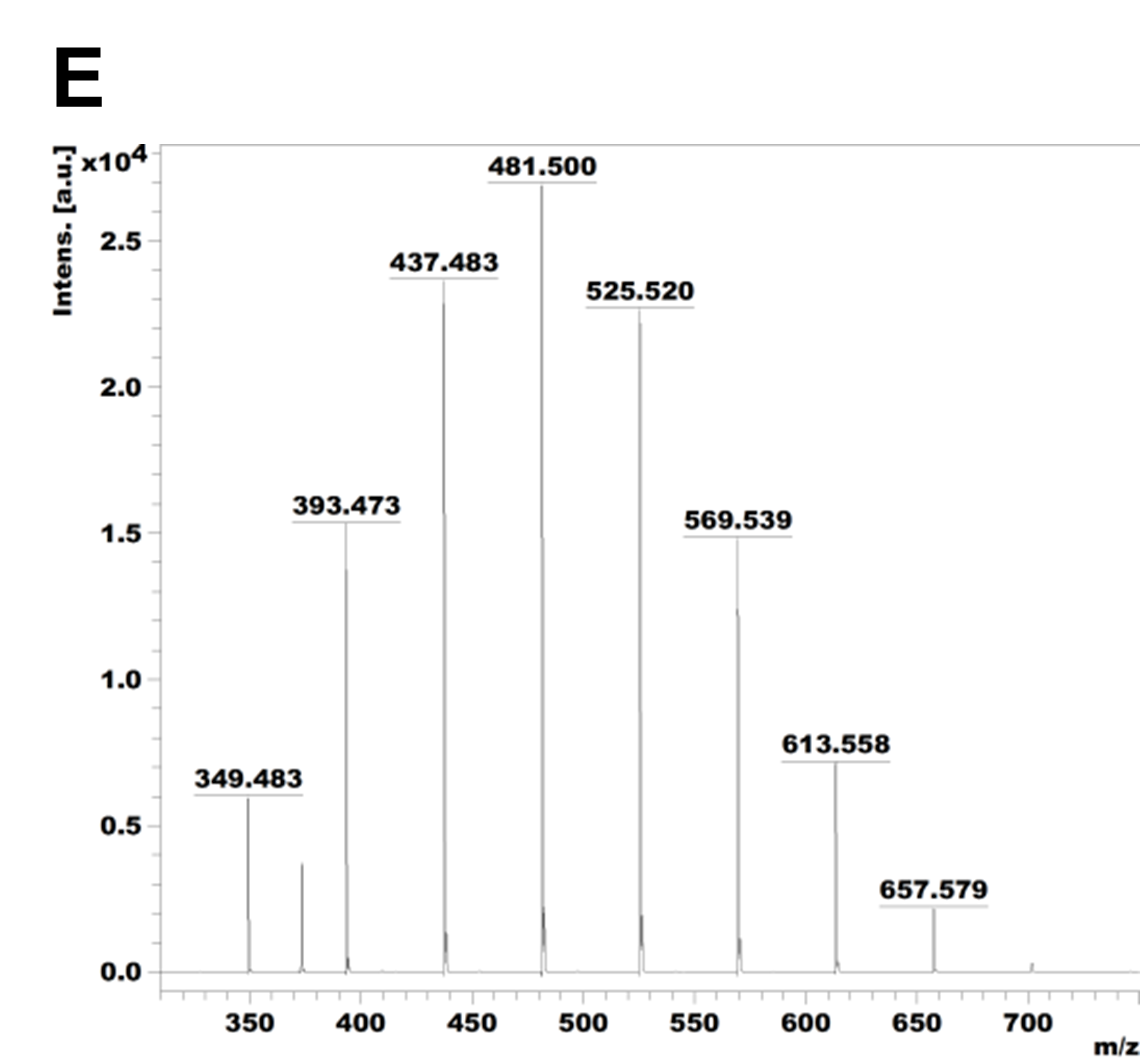

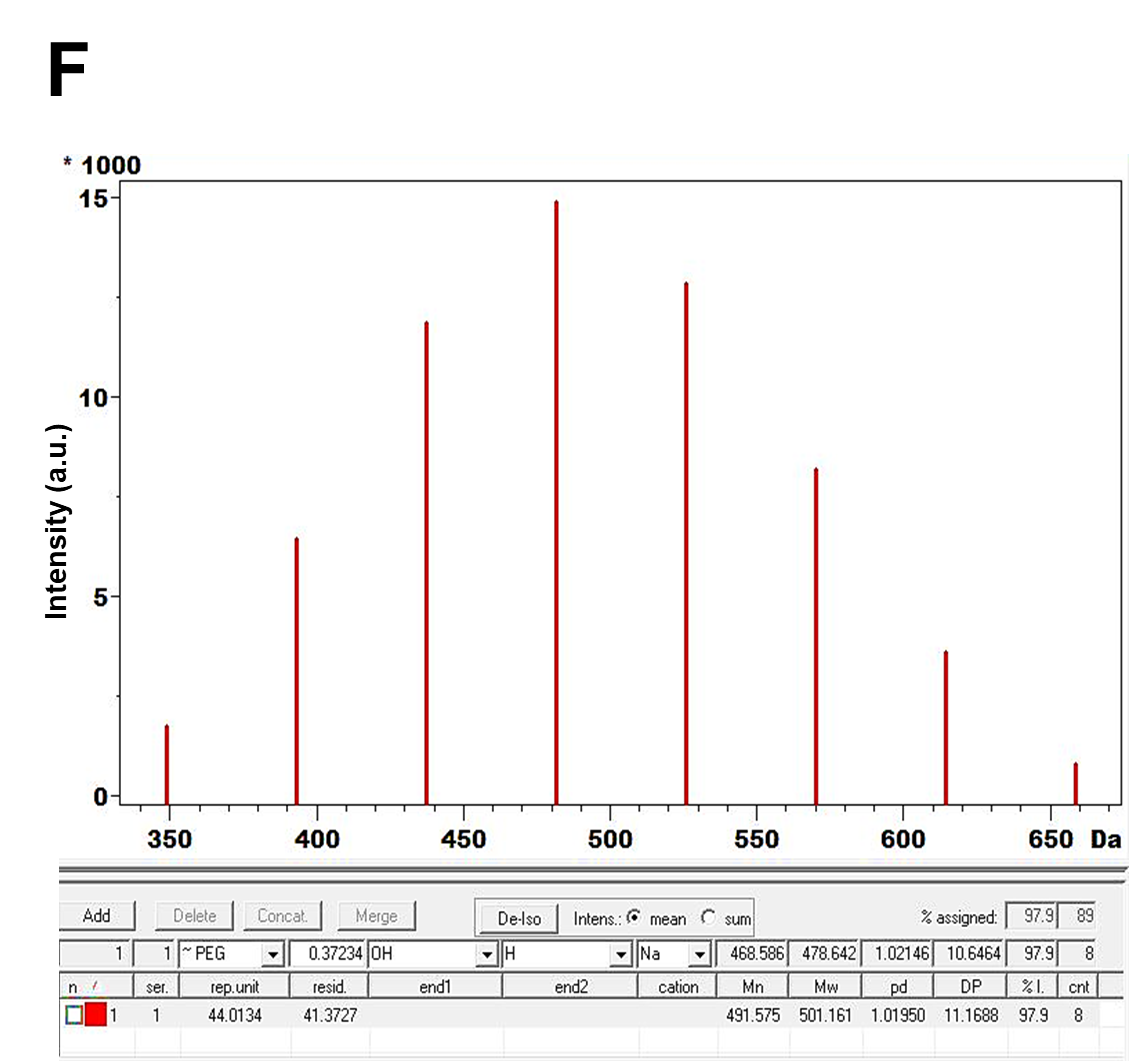

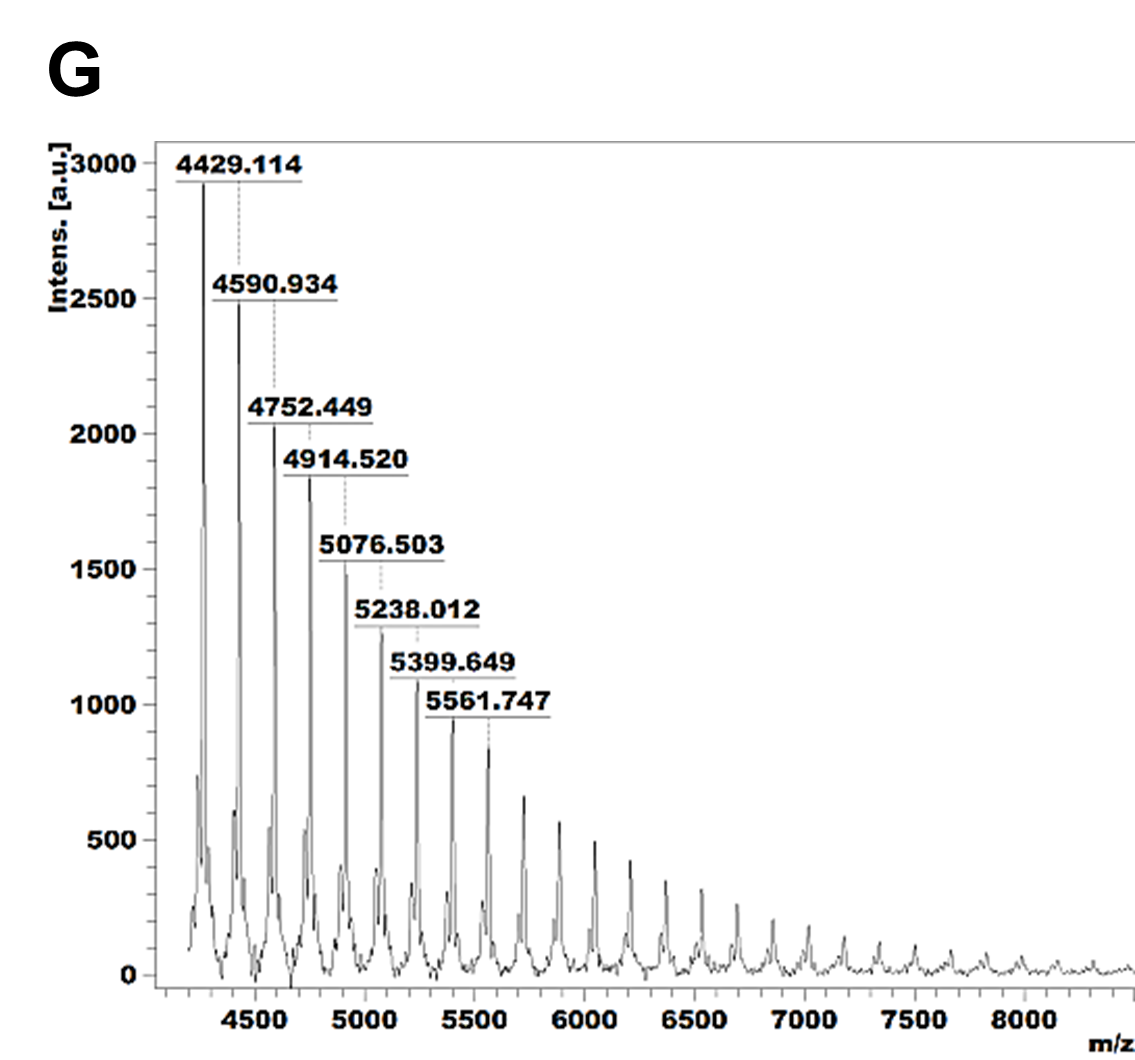

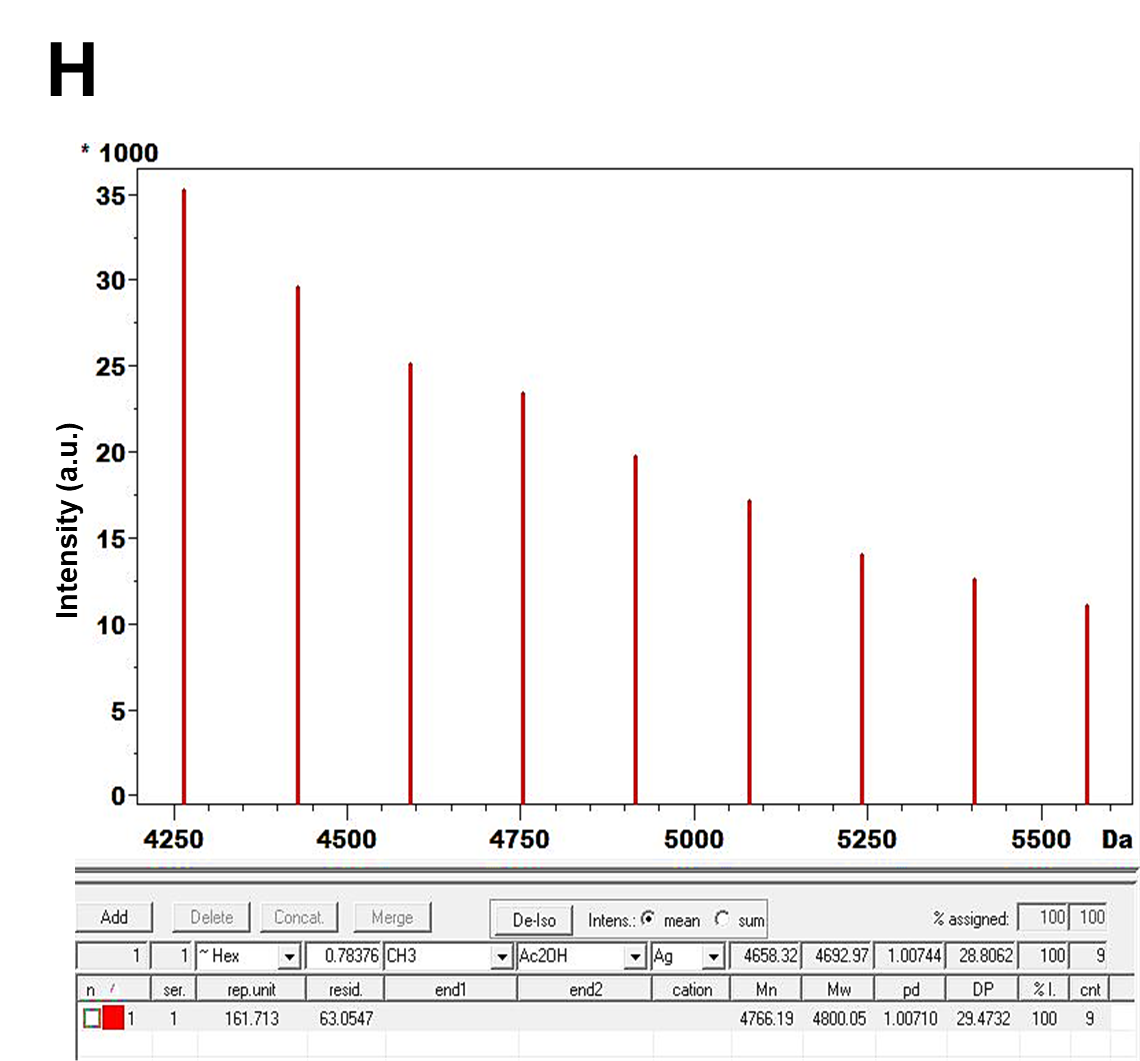

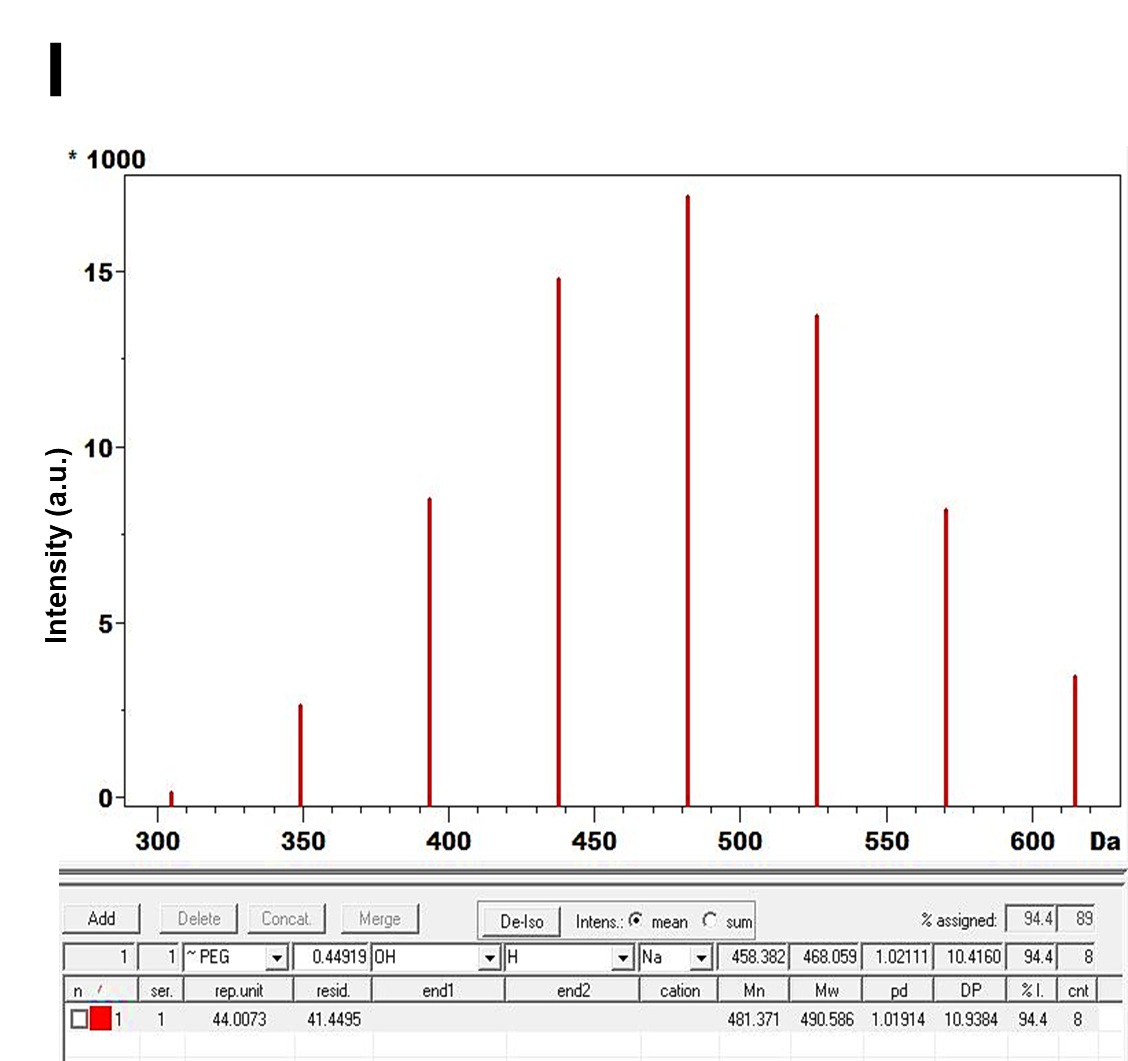

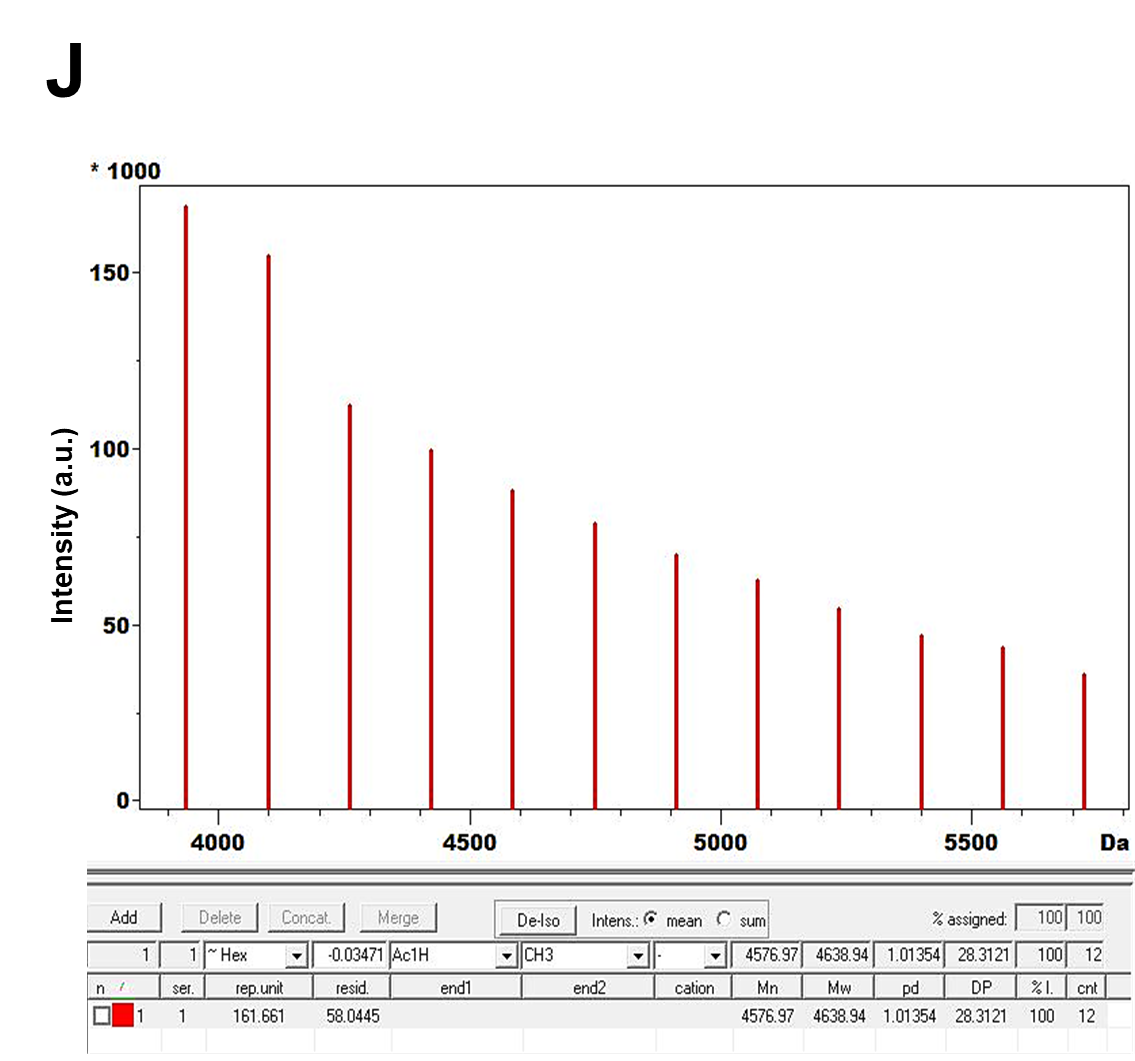

**Figure S6. MALDI-TOF mass spectrometric characterization of PEG 400–Dextran hydrogels and their components.** Matrix-assisted laser desorption/ionization time-of-flight (MALDI-TOF) spectra were recorded to identify polymeric constituents and confirm phase-specific polymer distribution in PEG 400–dextran (DEX) systems. **(A-B)** PEG 400 shows characteristic peaks separated by ~44 Da intervals, corresponding to ethylene oxide repeating units, as verified by PolyTools analysis, **(C-D)** Spectra of the supernatant from PEG 400/DEX 70 kDa hydrogels display similar ~44 Da spacing, indicating the presence of PEG in the soluble phase, **(E-F)** Supernatants from PEG 400/DEX 500 kDa gels exhibit comparable ~44 Da intervals, further confirming PEG enrichment in the supernatant, **(G-H)** The DEX 10 kDa spectrum displays peaks separated by ~162 Da intervals, characteristic of D-glucose repeating units as verified by PolyTools analysis, **(I)** PolyTools analysis of the supernatant from PEG 400/DEX 10 kDa gels exhibit comparable ~44 Da intervals, further confirming PEG enrichment in the supernatant, **(J)** PolyTools analysis of PEG400/DEX 10kDa hydrogel reveals ~162 Da spacing, confirming dextran as the primary constituent of the condensed phase. Together, these spectra demonstrate that PEG predominantly localizes in the supernatant, while dextran forms the condensed gel-like fraction, consistent with their monomeric repeating units.

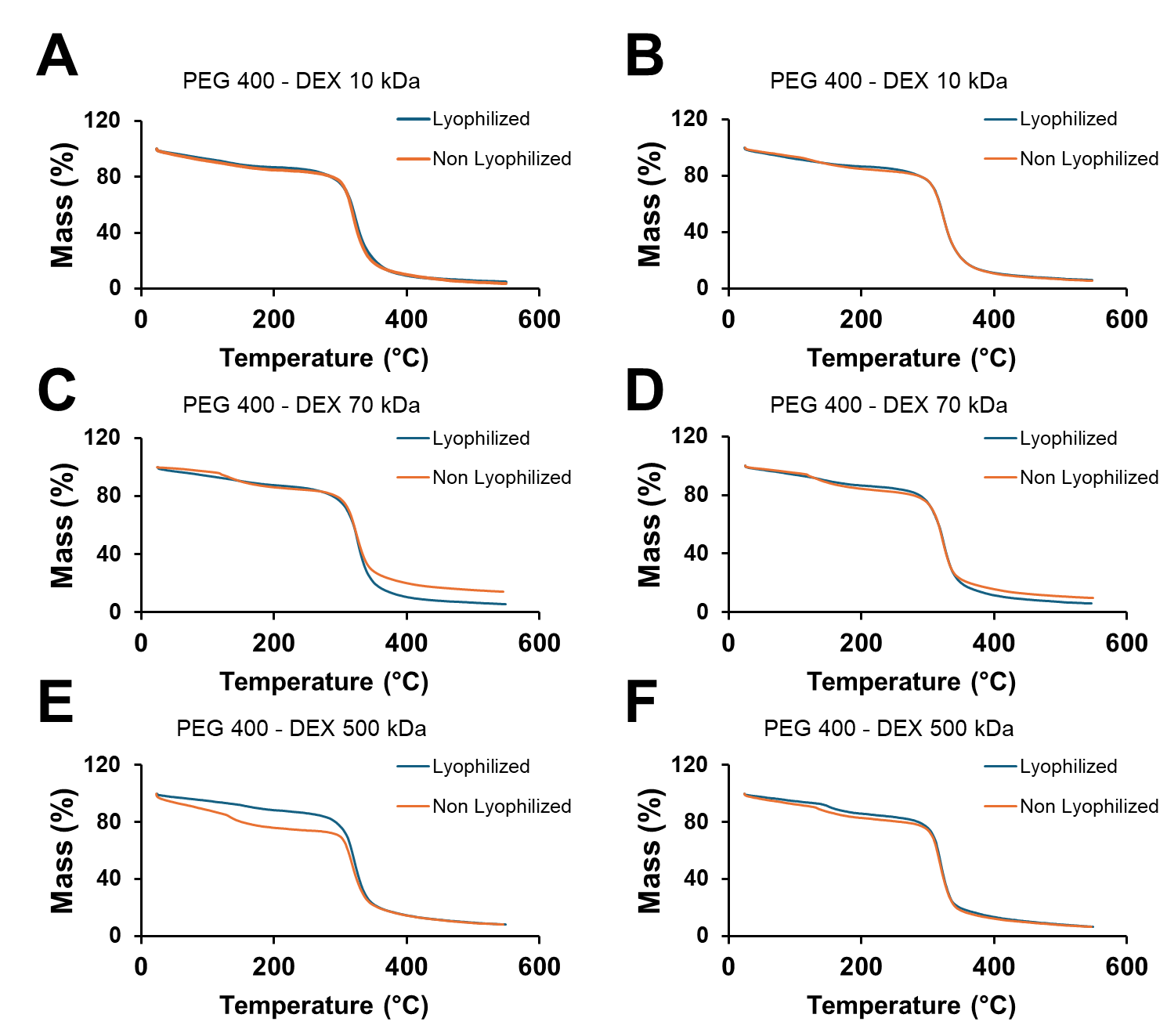

**Figure S7. Thermogravimetric analysis (TGA) of PEG 400 – DEX hydrogels.** TGA was performed to investigate solvent entrapment in lyophilized and non-lyophilized PEG 400–dextran (DEX) hydrogels. Formulations containing dextran of varying molecular weights **(A-B)** 10 kDa, **(C-D)** 70 kDa, and **(E-F)** 500 kDa were examined to identify the presence of entrapped solvent within the polymer matrices.

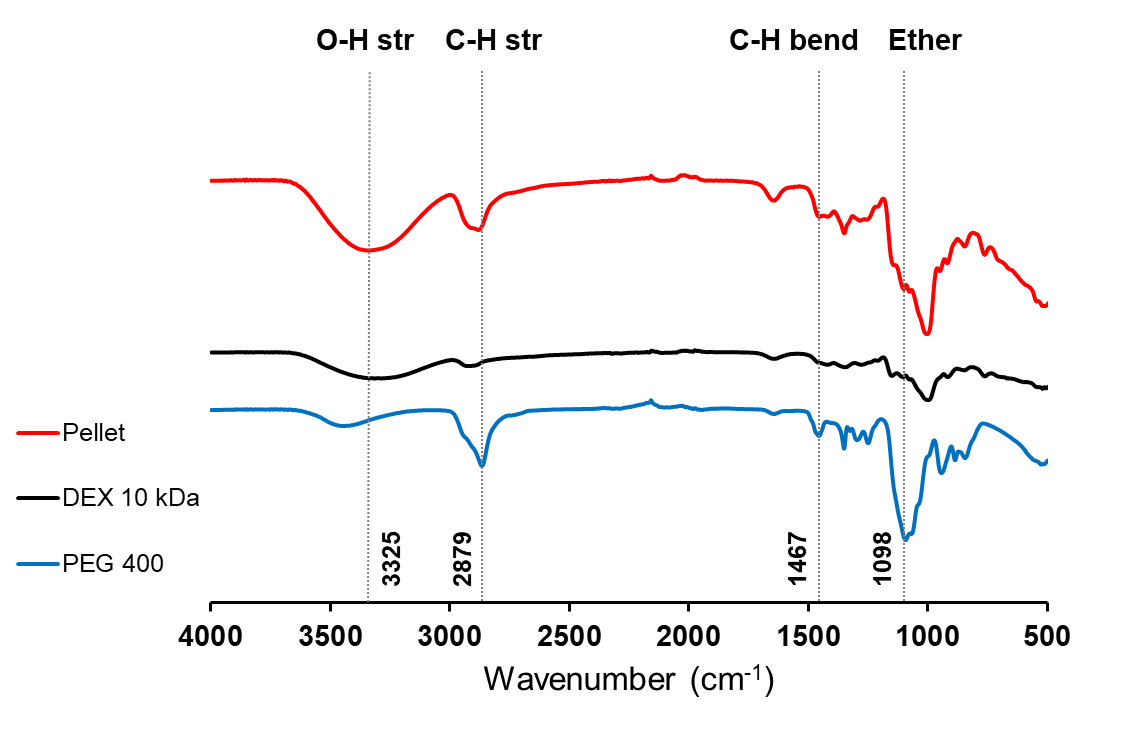

**Figure S8. Fourier Transform Infrared (FTIR) Spectra.** The FTIR spectra of PEG 400, DEX 10 kDa, and the hydrogel showed characteristic bands of both PEG and DEX in the hydrogel, showing the presence of contributions from each polymer component.

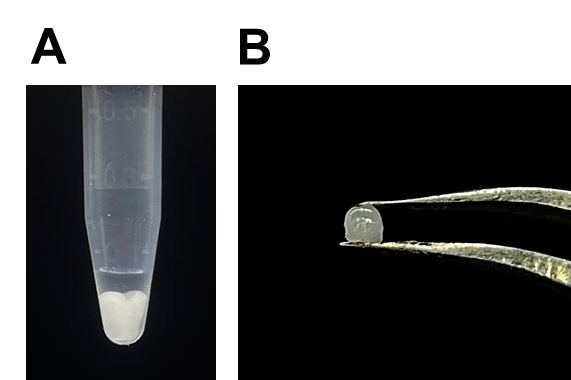

**Figure S9. Formation of Hydrogel with Poly(2-ethyl-2-oxazoline) (PEOx) and DEX 70 kDa. (A)** Soft hydrogel is formed when 100% PEOx and 20% DEX 70 kDa were mixed and vortexed, **(B)** Soft hydrogel when removed from the microcentrifuge tube, showing easy handleability.
